# Meta-analysis of 2,104 trios provides support for 10 novel candidate genes for intellectual disability

**DOI:** 10.1101/052670

**Authors:** Stefan H. Lelieveld, Margot R.F. Reijnders, Rolph Pfundt, Helger G. Yntema, Erik-Jan Kamsteeg, Petra de Vries, Bert B.A. de Vries, Marjolein H. Willemsen, Tjitske Kleefstra, Katharina Löhner, Maaike Vreeburg, Servi Stevens, Ineke van der Burgt, Ernie M.H.F. Bongers, Alexander P.A. Stegmann, Patrick Rump, Tuula Rinne, Marcel R. Nelen, Joris A. Veltman, Lisenka E.L.M. Vissers, Han G. Brunner, Christian Gilissen

**Affiliations:** Department of Human Genetics, Radboud Institute for Molecular Life Sciences, Radboud University Medical Center, Geert Grooteplein 10, 6525 GA Nijmegen, the Netherlands; Department of Human Genetics, Donders Centre for Neuroscience, Radboudumc, Geert Grooteplein 10, 6525 GA Nijmegen, the Netherlands; Department of Clinical Genetics, Maastricht University Medical Centre, Universiteitssingel 50, 6229 ER Maastricht, the Netherlands; Department of Genetics, University Medical Center Groningen, Hanzeplein 1, 9713 GZ Groningen, the Netherlands.

## Abstract

To identify novel candidate intellectual disability genes, we performed a meta-analysis on 2,637 *de novo* mutations, identified from the exomes of 2,104 ID trios. Statistical analyses identified 10 novel candidate ID genes, including *DLG4, PPM1D, RAC1, SMAD6, SON, SOX5, SYNCRIP, TCF20, TLK2* and *TRIP12*. In addition, we show that these genes are intolerant to non-synonymous variation, and that mutations in these genes are associated with specific clinical ID phenotypes.

## MAIN TEXT

Intellectual disability (ID) and other neurodevelopmental disorders are in part due to *de novo* mutations affecting protein-coding genes.^1^–^4^ Large scale exome sequencing studies of patient-parent trios have efficiently identified genes enriched for *de novo* mutations in cohorts of individuals with ID compared to controls,^2^ or based on expected gene-specific mutation rates.^5^

Here we sequenced the exomes of 820 patients with intellectual disability and their parents as part of routine genetic testing at the RadboudUMC in the Netherlands. We identified 1,083 *de novo* mutations (DNMs) in the coding and canonical splice site regions affecting 915 genes (**Supplementary Tables 1**–**2**, **Supplementary Figures 1**–**4**). In our cohort we detected an increased number of Loss-of-function (LoF) mutations compared to controls (Fisher’s exact test p=9.38x10^−12^, **Supplementary Methods**), and enrichment for recurrent gene mutations (observed vs. expected, p<1×10^−5^, **Supplementary Figure 5**).

Based on an established framework of gene specific mutation rates^6^, we calculated for each gene the probability of identifying the observed number of LoF or functional DNMs in our cohort (**Supplementary Methods**). To validate this approach we first performed the analysis on the complete set of 820 ID patients. After Benjamini-Hochberg correction for multiple testing, 18 well-known ID genes were significantly enriched for DNMs (**Supplementary Table 3** and **4**). To optimize our analysis for the identification of novel candidate genes in the RUMC cohort, we removed all individuals with mutations in any of the known ID genes (**Supplementary Methods**, **Supplementary Figure 6**). Repeating the analysis for mutation enrichment, we identified 4 genes (*DLG4*, *PPM1D*, *SOX5*, *TCF20*) not previously associated with ID to be significantly enriched for DNMs in our cohort (**Figure 1**, **Table 1**, **Supplementary Table 5**). To achieve the best possible power for the identification of novel candidate ID genes, we next added data from 4 previously published family-based sequencing studies (**Supplementary Table 1**). The combined cohort included 2,104 trios and 2,637 DNMs across 1,990 genes. After again excluding individuals with mutations in known ID genes, this cohort consisted of 1,471 individuals with 1,400 DNMs in 1,235 genes (**Supplementary Methods**, **Supplementary Figure 6**). Meta-analysis on this combined cohort identified 10 novel candidate ID genes with more LoF, or more functional, DNMs than expected *a priori*. These 10 genes included the four novel candidate ID genes previously identified in the RUMC cohort, as well as *RAC1*, *SMAD6*, *SON*, *TLK2*, *TRIP12* and *SYNCRIP* (**Figure 1**, **Table 1**, **Supplementary Table 6**).

**Figure 1.**
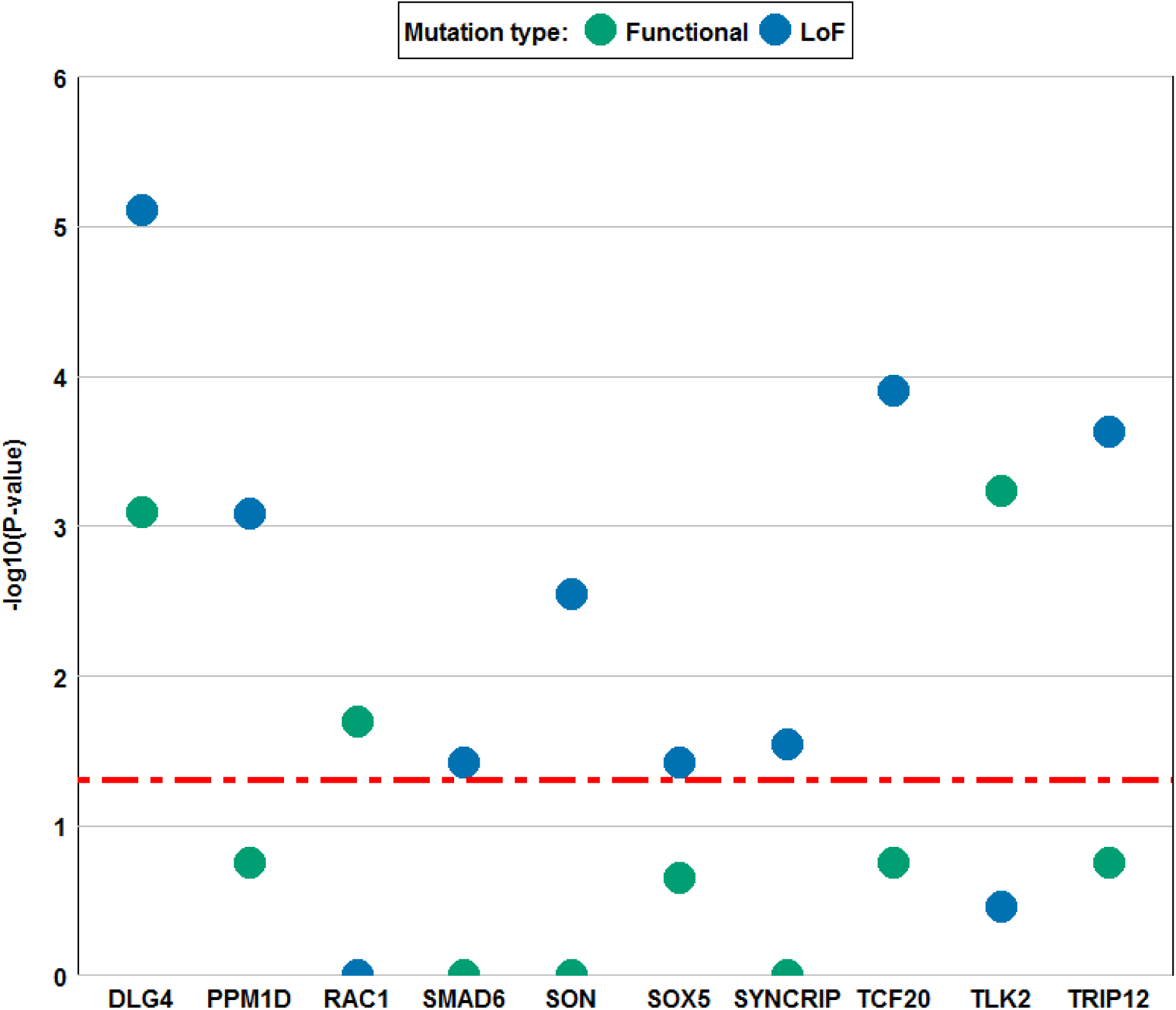
Genes enriched for LoF and functional *de novo* mutations (DNMs) in a cohort of 2,104 ID trios from multiple studies. The y-axes shows the-log10 transformed corrected P-value of the *de novo* mutation enrichment. Corrected P-values based on LoF mutations are colored in blue and corrected P-values based on functional mutations are colored green. Only genes with a corrected P-value (LoF, functional, or both) less than the significance threshold (red dotted line, 0.05) are shown.

**Table 1.**
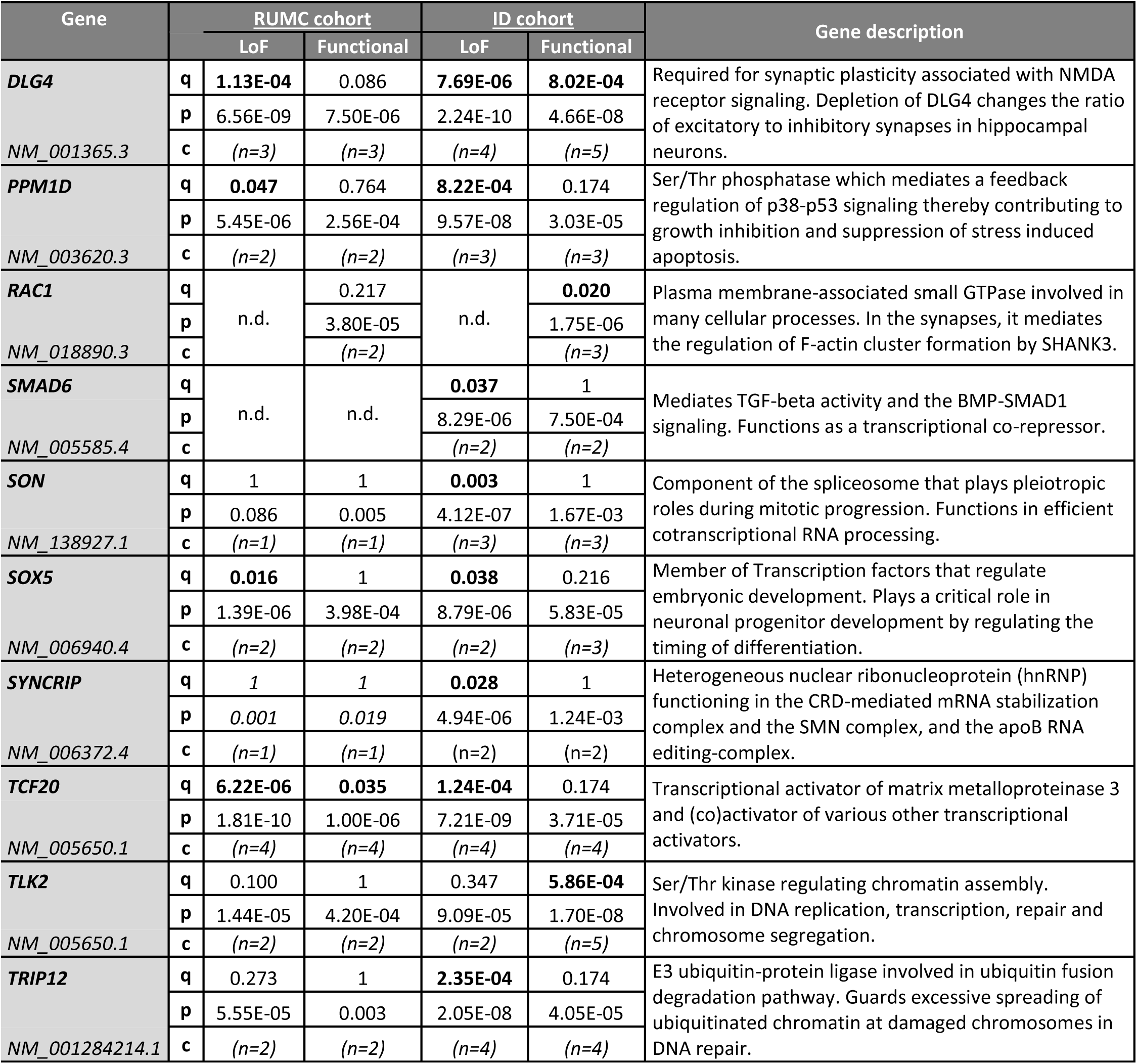
Novel candidate ID genes. All genes listed reached statistical significance after Benjamini-Hochberg correction for enrichment of functional and/or loss-of-function (LoF) DNM in the RUMC or ID cohort. For each gene the Benjamini-Hochberg corrected p-value (**q**), uncorrected p-value (**p**) and the raw counts (**c**) are shown.

| Gene | RUMC cohort |  |  | ID cohort |  | Gene description |
| --- | --- | --- | --- | --- | --- | --- |
|  |  | LoF | Functional | LoF | Functional |  |
| **DLG4**<br>NM_001365.3 | **q** | **1.13E-04** | 0.086 | **7.69E-06** | **8.02E-04** | Required for synaptic plasticity associated with NMDA receptor signaling. Depletion of DLG4 changes the ratio of excitatory to inhibitory synapses in hippocampal neurons. |
|  | **p** | 6.56E-09 | 7.50E-06 | 2.24E-10 | 4.66E-08 |  |
|  | **c** | (n=3) | (n=3) | (n=4) | (n=5) |  |
| **PPM1D**<br>NM_003620.3 | **q** | **0.047** | 0.764 | **8.22E-04** | 0.174 | Ser/Thr phosphatase which mediates a feedback regulation of p38-p53 signaling thereby contributing to growth inhibition and suppression of stress induced apoptosis. |
|  | **p** | 5.45E-06 | 2.56E-04 | 9.57E-08 | 3.03E-05 |  |
|  | **c** | (n=2) | (n=2) | (n=3) | (n=3) |  |
| **RAC1**<br>NM_018890.3 | **q** | n.d. | 0.217 | n.d. | **0.020** | Plasma membrane-associated small GTPase involved in many cellular processes. In the synapses, it mediates the regulation of F-actin cluster formation by SHANK3. |
|  | **p** |  | 3.80E-05 |  | 1.75E-06 |  |
|  | **c** |  | (n=2) |  | (n=3) |  |
| **SMAD6**<br>NM_005585.4 | **q** | n.d. | n.d. | **0.037** | 1 | Mediates TGF-beta activity and the BMP-SMAD1 signaling. Functions as a transcriptional co-repressor. |
|  | **p** |  |  | 8.29E-06 | 7.50E-04 |  |
|  | **c** |  |  | (n=2) | (n=2) |  |
| **SON**<br>NM_138927.1 | **q** | 1 | 1 | **0.003** | 1 | Component of the spliceosome that plays pleiotropic roles during mitotic progression. Functions in efficient cotranscriptional RNA processing. |
|  | **p** | 0.086 | 0.005 | 4.12E-07 | 1.67E-03 |  |
|  | **c** | (n=1) | (n=1) | (n=3) | (n=3) |  |
| **SOX5**<br>NM_006940.4 | **q** | **0.016** | 1 | **0.038** | 0.216 | Member of Transcription factors that regulate embryonic development. Plays a critical role in neuronal progenitor development by regulating the timing of differentiation. |
|  | **p** | 1.39E-06 | 3.98E-04 | 8.79E-06 | 5.83E-05 |  |
|  | **c** | (n=2) | (n=2) | (n=2) | (n=3) |  |
| **SYNCRIP**<br>NM_006372.4 | **q** | 1 | 1 | **0.028** | 1 | Heterogeneous nuclear ribonucleoprotein (hnRNP) functioning in the CRD-mediated mRNA stabilization complex and the SMN complex, and the apoB RNA editing-complex. |
|  | **p** | 0.001 | 0.019 | 4.94E-06 | 1.24E-03 |  |
|  | **c** | (n=1) | (n=1) | (n=2) | (n=2) |  |
| **TCF20**<br>NM_005650.1 | **q** | **6.22E-06** | **0.035** | **1.24E-04** | 0.174 | Transcriptional activator of matrix metalloproteinase 3 and (co)activator of various other transcriptional activators. |
|  | **p** | 1.81E-10 | 1.00E-06 | 7.21E-09 | 3.71E-05 |  |
|  | **c** | (n=4) | (n=4) | (n=4) | (n=4) |  |
| **TLK2**<br>NM_005650.1 | **q** | 0.100 | 1 | 0.347 | **5.86E-04** | Ser/Thr kinase regulating chromatin assembly. Involved in DNA replication, transcription, repair and chromosome segregation. |
|  | **p** | 1.44E-05 | 4.20E-04 | 9.09E-05 | 1.70E-08 |  |
|  | **c** | (n=2) | (n=2) | (n=2) | (n=5) |  |
| **TRIP12**<br>NM_001284214.1 | **q** | 0.273 | 1 | **2.35E-04** | 0.174 | E3 ubiquitin-protein ligase involved in ubiquitin fusion degradation pathway. Guards excessive spreading of ubiquitinated chromatin at damaged chromosomes in DNA repair. |
|  | **p** | 5.55E-05 | 0.003 | 2.05E-08 | 4.05E-05 |  |
|  | **c** | (n=2) | (n=2) | (n=4) | (n=4) |  |

To further evaluate the identification of the 10 novel candidate ID genes, we compared the phenotypes of the 18 RUMC individuals with DNMs in these genes. We observed strong phenotypic overlap for some of these genes (**Figure 2**, **Supplemental case reports**, **Supplementary Table 7**). Further genes, such as *SETD2*, which are close to statistical significance, show phenotypic similarities suggestive for a shared genetic cause consistent with previous case reports^7^, ^8^ (**Supplementary Figure 7**, **Supplemental Case reports**).

**Figure 2.**
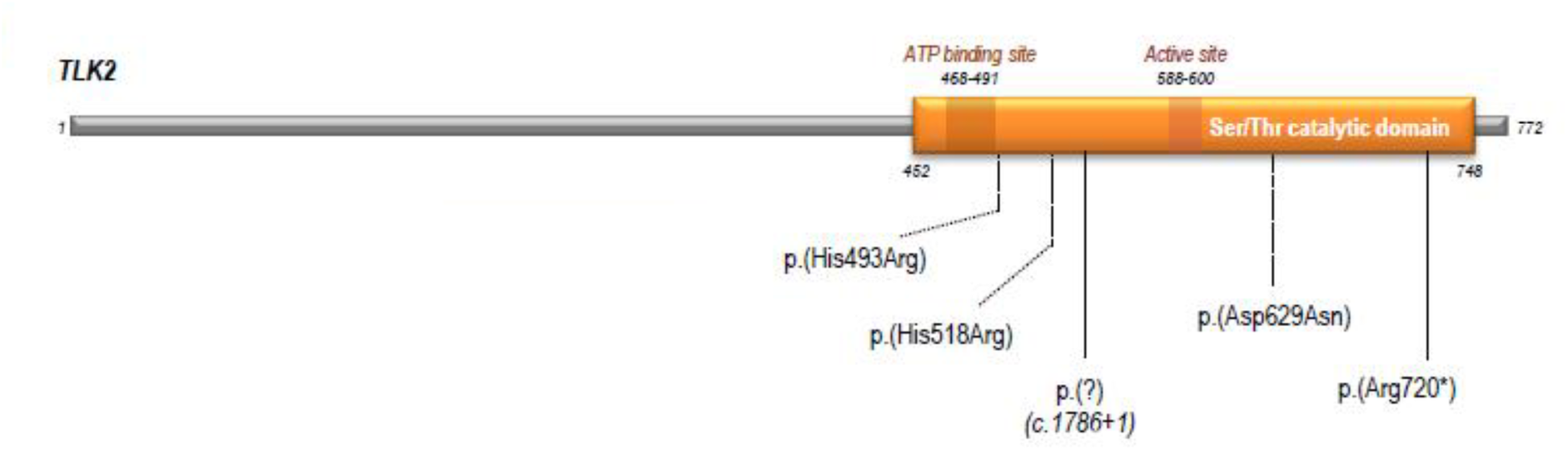
TLK2 protein (Q86UE8) with *de novo* mutations localized to the serine/threonine catalytic domain. Both individuals in the RUMC cohort with a DNM in *TLK2* show overlapping clinical features including facial dysmorphisms **(Supplemental Case reports)**.

Studies have shown that genes involved in genetic disorders exhibit strongly reduced tolerance to genetic non-synonymous variation compared to non-disease genes. This is particularly evident for ID.^3^ We found that a large set of well-known dominant ID genes (n=444), as well as the 10 novel candidate ID genes are highly intolerant to LoF variation^9^ (Median pLI of 0.95; p<1×10^−5^ and 0.99; p<1×10^−5^ respectively; **Supplementary Methods**, **Supplementary Figure 8** and **Supplementary Table 8**). Intriguingly, we noted that those known and novel candidate dominant ID genes that harbor only missense variants are among the most intolerant ID genes (Median pLI of 0.99; p<1×10^−5^; **Supplementary Figure 8**). Additionally, we find that mutations in ‘missense only’ genes are more likely to cluster than mutations in genes for which we also identified LoF mutations (p=0.01, Fisher’s exact test; **Supplementary Methods**, **Supplementary Table 9**).

There is considerable overlap of genes and molecular pathways involved in neurodevelopmental disorders (NDDs) such as autism spectrum disorder (ASD), schizophrenia (SCZ), epileptic encephalopathy (EE) and ID.^10^ Therefore we performed a third analysis including 12 published family-based sequencing studies of various NDDs (**Supplementary Table 1**,**Supplementary Figure 6**). Repeating our analysis in this NDD cohort, we identified 7 genes significantly enriched for either LoF or functional DNMs (**Supplementary Figure 9**, **Supplementary Table 10**). In line with our hypothesis, five of the identified genes were also identified in our previous analyses with individuals with ID only, whereas two genes (*SLC6A1* and *TCF7L2*) only reached significance in the NDD meta-analysis due to additional mutations in patients with other phenotypes than ID (**Supplementary Table 11**). In more detail, for two of the five candidate ID genes (*TLK2* and *TRIP12*) additional *de novo* mutations were identified in individuals with ASD and SCZ, suggesting that DNMs in these genes may lead to a broader phenotype than only ID. For *TRIP12*, a similarly broad phenotype has been reported previously.^4^

In summary, we identified 10 novel candidate ID genes in a meta-analysis of WES data on 2,104 ID trios. The statistical framework used here, differs from existing methods based on gene specific mutation rates, by removing all trios with mutations in known disease genes, and by applying Benjamini-Hochberg correction for multiple testing. Our study underscores the impact of *de novo* mutations on a continuum of neurodevelopmental phenotypes, that impinge on a broad range of processes including chromatin modifiers (*TRIP12*, *TLK2*), FMRP target and synaptic plasticity genes (*DLG4;* **Supplementary Figure 10**) and embryonically expressed genes (*PPM1D*, *RAC1*)^2^. Data from a similar systematic study of *de novo* mutations in neurodevelopmental disorders suggest that many, and possibly most, genes causing severe developmental disorders by *de novo* mutation are now known^11^. Yet, only *TCF20* and *PPM1D* are shared between the 10 novel genes in our study and the 14 genes identified by McRae and colleagues. Thus, a large number of rare dominant developmental disorder genes may remain to be identified.

## Contributions

C.G., L.E.L.M.V. and H.G.B. designed the study; S.H.L., M.R.F.R., C.G., L.E.L.M.V., performed the analysis. R.P., H.G.Y., E.K., T.R., S.S., A.P.A.S. and M.R.N. signed out initial diagnosed reports. P.dV. performed Sanger validations. B.B.A.dV., M.H.W., T.K., K.L., M.V., I.vdB., E.M.H.F.B., P.R. and M.R.F.R., collected patient phenotypes. S.H.L., M.R.F.R., J.A.V., H.G.B., L.E.L.M.V. and C.G., drafted the manuscript, all authors contributed to the final version of the paper.

## Online Methods

### Recruitment of individuals with ID

The Department of Human Genetics from the Radboud University Medical Center (RUMC) is a tertiary referral center for clinical genetics. Approximately 350 individuals with unexplained intellectual disability (ID) are referred annually to our clinic for diagnostic evaluation. Since September 2011 whole exome sequencing (WES) is part the routine diagnostic work-up aimed at the identification of the genetic cause underlying disease.^12^ For individuals with unexplained ID, a family-based WES approach is used which allows the identification of *de novo* mutations as well as variants segregating according to other types of inheritance, including recessive mutations and maternally inherited X-linked recessive mutations in males.^13^ For the purpose of this study, we selected all individuals with ID who had family-based WES using the Agilent SureSelect v4 enrichment kit combined with sequencing on the Illumina HiSeq platform in the time period 2013-2015. This selection yielded a set of 820 individuals, including 359 females and 461 males. The level of ID ranged between mild (IQ 50-70) and severe-profound (IQ<30).

Families gave informed consent for both the diagnostic procedure as well as for forthcoming research that could result in the identification of new genes underlying ID by meta-analysis, as presented here. Explicit consent for photo-publication was sought and given by a subset of families.

### Diagnostic whole exome sequencing

The exomes of 820 patient-parent trios were sequenced using DNA isolated from blood, at the *Beijing Genomics Institute* (BGI) in Copenhagen. Exome capture was performed using Agilent SureSelect v4 and samples were sequenced on an Illumina HiSeq instrument with 101bp paired-end reads to a median coverage of 75x. Sequence reads were aligned to the hg19 reference genome using BWA version 0.5.9-r16. Variants were subsequently called by the GATK unified genotyper (version 3.2-2) and annotated using a custom diagnostic annotation pipeline. Base-pair resolution coverage of the regions enriched by the SureSelect V4 kit are computed by BEDTools based on the regions as provided by the manufacturer. An average 98.9% of Agilent SureSelect V4 enriched targets was covered by 10 or more reads for the RUMC cohort of 820 ID patients (**Supplementary Figure 1**).

### Identification of *de novo* mutations in 820 individuals with ID

The diagnostic WES process as outlined above only reports (*de novo*) variants that can be linked to the individuals’ phenotype. In this study, we systematically collected all *de novo* mutations located in coding sequence (RefSeq) and/or affected canonical splice sites (canonical dinucleotides GT and AG for donor and acceptor sites; **Supplementary Figure 4**), identified in 820 individuals with ID irrespective of their link to disease, to evaluate the potential relevance of genes for ID in an unbiased fashion using a statistical framework. *De novo* mutations were called as described previously.^13^ Briefly, variants called within parental samples were removed from the variants called in the child. For the remaining variants pileups were generated from the alignments of the child and both parents. Based on pileup results variants were then classified into the following categories: “maternal (for identified in the mother only)”, “paternal (for identified in the father only)”, “low coverage” (for insufficient read depth in either parent), “shared” (for identified in both parents)”, and “possibly *de novo*” (for absent in the parents). Variants classified as “possibly *de novo*” were included in this study.

We applied various quality measures to ensure that only the most reliable calls were included in the study: (i) all samples had less than 25 “possibly *de novo*” calls; (ii) the variant had at least 10x coverage in either parent (e.g. high prior probability of being inherited); (iii) the location was not known in dbSNP version137 (e.g. possible highly mutable genomic location) and (iv) was identified in a maximum of 5 samples in our in-house variant database (e.g. eliminating variants that are too frequent to be disease-causing given the incidence of ID in combination with the sample size of our in-house database); and (v) each variant showed a variant read percentage >30%, or alternatively, >20% with >10 individual variant reads and (vi) a GATK quality score of >400. For *de novo* variants called within a 5bp window of each other within the same individual, variant calls were manually curated and merged into a single call (when occurring on the same allele). This set of criteria resulted in the identification of 1,083 potential *de novo* mutations in 820 individuals with ID.

### Validation and categorization of *de novo* mutations

In a separate (unpublished) in-house study, we recently determined the predictive value for GATK quality scores in terms of the variant being validated by Sanger sequencing. A set of 840 variants called by the same version of GATK was retrospectively analyzed for their respective quality scores and validation status. Based on this assessment, we determined that a GATK quality score ?500 resulted in 100% of variants being validated by Sanger sequencing (data not shown; internal reference VAL G 084). In addition to our in-house study two other studies also found a 100% Sanger validation rate for the variants with a GATK quality score of ≥500.^14^ Based on this we considered all variants with a GATK Q-score of ≥500 (n=1,039) to be true *de novo* mutations. Nonetheless, a random set of 141 potential *de novo* mutations with GATK Q-scores of ≥500 were all confirmed by Sanger sequencing. All potential *de novo* variants with a GATK Q-score between 400-500 (n=40) were subsequently validated by Sanger sequencing, and all were confirmed. All 20 *de novo* mutations of the reported novel candidate genes were confirmed by Sanger sequencing (**Supplementary Table 2**, **Supplementary Figures 2**–**4**).

For further downstream statistical analysis (see below), DNMs were categorized by mutation type: (i) **loss-of-function (LoF) DNM (n=211)**, including nonsense (n=77), frameshift (n=97), canonical splice site (n=27), start loss (n=2), stop loss (n=1), premature stop codon resulting from an indel (n=7); (ii) **functional DNM (n=872)**, including all LoF mutations (n=211), in-frame insertion/deletion events (n=23) and all missense mutations (n=638) (**Supplementary Figure 4**). For variants within the same individual and within the same gene but more than 5bp apart, the variant with the most severe functional effect was considered for the per gene statistics (see below).

### Evaluating the number of recurrently LoF and functional *de novo* mutated genes

We simulated the expected number of recurrently mutated genes by redistributing the observed number of mutations at random over all genes based on their specific loss-of-function (LoF, see section “Statistical enrichment of DNMs”) and functional mutation rates (see section “Statistical enrichment of DNMs”) as described by Samocha *et al.*^6^ Based on 100,000 simulations we calculated how many times the number of recurrently mutated genes was the same or exceeded the observed number of recurrently mutated genes in the RUMC dataset. We performed the simulation separately for LoF and functional DNMs (**Supplementary Figure 5**). P-values were then calculated by taking the number of times the number of recurrently mutated genes exceeded the observed number of recurrently mutated genes, divided by the number of simulations.

### Genes previously implicated in ID etiology

To evaluate whether the genes identified by our meta-analyses have been implicated in ID before, two publicly available repositories of genes known to be involved in ID were used. Firstly, we used our list of 707 genes, routinely used by our diagnostic setting to interpret WES results of individuals with ID.^15^ Secondly, we downloaded a list of 1,424 genes from the DDG2P database reflecting genes to be associated with developmental disorders, which has been compiled and curated by clinicians as part of the DDD study to facilitate clinical feedback of likely causal variants.^5^ In total the two lists compromised 1,537 unique genes. In this manuscript, the list of unique gene entries is referred to as known ID genes (**Supplementary Table 4**).

### Statistical enrichment of DNMs

In our meta-analysis for ID and neurodevelopmental disorders we only included studies with minimum of 50 trios. For each gene, and each of the functional classes (LoF and functional), we used the corresponding gene specific mutation rate (GSMR) as published by Samocha *et al.* (2014)^6^ to calculate the probability of the number of identified *de novo* mutations in our cohort. For genes for which no GSMR was reported, we used the maximum GSMR of all reported genes (*i.e*. the GSMR of the gene *TTN*). We then calculated specific mutation rates for the two defined functional classes (loss-of-function, functional). The GSMR for loss-of-function DNMs was calculated by summing the individual GSMR for nonsense, splice site and frame-shift variants; The GSMR for functional DNMs was calculated by summing the GSMR for the loss-of-function (LoF) variants with the missense mutation rate; and for genes for which variants from different functional classes were identified, we used the overall GSMR. For the stoploss and startloss mutations we used the LoF-rate and for in-frame indels the functional rate. Null hypothesis testing was done using a one-sided exact Poisson Test based on a sample size of 820 individuals with ID, representing 1,640 alleles for autosomal genes, and 1,179 alleles for genes on the X-chromosome (461 males).

For DNMs on chromosome X the correct mutation rate depends on the patient’s gender as the mutation rates for fathers is higher than for mothers. Estimates show a 4:1 ratio of paternal to maternal *de novo* mutations^16^. Hence, male offspring, receiving their chromosome X exclusively from the mother, have therefore a lower mutation rate on chromosome X than estimated by the GSMR. This correction could however only be performed for the RUMC cohort, as information of gender was not available for all studies included in the ID cohort. Of note, not correcting for this bias in male individuals for DNM in genes on the X chromosome, however will lead to less significant p-values for genes on the X-chromosome, thereby potentially underestimating the significance of candidate novel ID genes located on the X-chromosome. In the case where one patient has two DNMs in the same gene we ignored one of the two DNMs for the statistical enrichment analysis to avoid false positive results. In the aforementioned case the severity of the DNM protein-effect was used in the choice which DNM to ignore. For example, if a patient has one missense and one nonsense DNM in the same gene the missense mutation was ignored in the statistical analysis.

The gene specific *p*-values were corrected for multiple testing based on the 18,730 genes (present in the Agilent V4 exome enrichment kit) times the number of tests (x2), using the Benjamini-Hochberg procedure with an FDR of 0.05. In our cohort of 820 individuals with ID, conclusive diagnosis were already made based on DNMs in a genes previously implicated in disease. The use of a multiple testing correction with a FDR of 0.05, in combination with a potential large number of DNMs in known ID genes may cause the artificial significance of other genes because of an increasing lenient correction for the least significant genes^17^. To verify that the identification of novel candidate ID genes is not inflated by this effect, we performed the analysis after removing all individuals with a DNM in one of the known genes (potential other DNMs in such individuals were also removed for further analysis). Incidentally, this also increased our statistical power. The mode of inheritance was not taken into account when removing individuals with a DNM in a known gene (e.g. samples with a DNM in a recessive gene were excluded). This correction left 584/820 individuals with ID in the RUMC cohort, with 627 DNMs across 584 genes. Similarly, for the ID and neurodevelopmental cohort, we removed all individuals with a DNM in a known ID gene (and other DNMs in these individuals). For the ID cohort, 1,471 samples remained with 1,400 DNMs in 1,235 genes. For the neurodevelopmental cohort, 4,944 samples with 4,387 DNM across 3,402 genes remained (For the complete overview see **Supplementary Figure 6**). We corrected for testing 34,386 genes (*i.e*. all 18,730 genes minus the 1,537 known ID genes multiplied by two for testing the loss-of-function and functional categories).

### Validation of the statistical approach by analysis of DNMs in a control cohort

To further confirm the validity of our statistical approach, we applied the same analyses to a set of DNMs identified in trios of healthy individuals and unaffected siblings. Hereto, we downloaded and re-annotated all DNMs identified in 1,911 unaffected siblings of individuals with ASD from Iossifov *et al.*^2^ together with DNMs in controls (**Supplementary Table 14**). In total the control set contained 2,019 coding DNMs found in 2,299 trios. Of note, the protein coding *de novo* mutation rate in the control cohort was markedly lower than observed in the individuals with ID (0.91 vs. 1.32, respectively). Additionally, we observed no significant enrichment of recurrently mutated genes for loss-of-function or functional mutations (p=0.60 and p=0.12, respectively, **Supplementary Figure 11**).

For the control cohort we performed the statistical analysis as described above and identified only one gene to be significantly enriched for DNMs. For *YIF1A* (FDR corrected p-value = 0.01) we identified a total of 3 missense and 1 frame-shift DNM (**Supplementary Table 13**–**14**). *YIF1A* may be involved in transport between the endoplasmic reticulum and the Golgi, and has a pLI (probability of being loss-of-function intolerant) of 2.08x×10^−8^ indicating this gene is a loss-of-function tolerant gene. We note that the control cohort consists mostly of healthy siblings from individuals with ASD, and, as such, may still have a small enrichment for mutations that lead to susceptibility for neurodevelopmental disorders.

### Increased number of Loss-of-function (LoF) mutations in RUMC cohort compared to controls

To reduce the impact of the used enrichment kit used in the control set studies and RUMC cohort we computed the intersection of all enrichment kits (Agilent SureSelect 37Mb Agilent SureSelect 50Mb Agilent SureSelect V4 Nimblegen SeqCap V2; see **Supplementary Table 12**) via the ´intersect´ function of BEDTools. Only the loss-of-function de novo mutations present in the 28,189,737 Mb intersection of the four enrichment kits were used in the analysis. The Fisher’s exact test on the enrichment kits normalized loss-of-function *de novo* mutations yielded a significant difference with p-value= 9.38×^−12^ (RUMC: 157 LoF *de novo* mutations of a total of 805 de novo mutations; control set: 137 LoF *de novo* mutations of a total of 1,485 de novo mutations). The coverage and other relevant technical information of the control studies are enlisted in **Supplementary table 12**. We note it is important to take into account the coverage and false negative rates of all sequencing studies. So far, only a single study has attempted to provide a false negative rate (e.g. mutations that are there but were not identified) for exome sequencing, and this was predicted to be <5%^2^.

### Attributing pLI for all protein coding genes

To determine the intolerance to loss-of-function (LoF) variation for each gene, we used the pLI which is based on data from the Exome Aggregation Consortium (ExAC) version 0.3.1 providing exome variants from 60,706 unrelated individuals^9^. The pLI, or the probability of being LoF intolerance, is based on the expected vs. observed variant counts to determine the probability that a gene is intolerant to LoF variants and is computed for a total of 18,226 genes. The closer a pLI is to 1 the more intolerant a gene is to LoF variants. The authors consider a pLI >= 0.9 as an extremely LoF intolerant set of genes. The list of the pLI for the genes used in this study can be found in **Supplementary Table 8**. The performance of the pLI was evaluated for four gene sets **1)** 170 Loss-of-function (LoF) tolerant genes^18^ **2)** 404 “House-keeping” genes, involved in crucial roles in cell maintenance^19^ **3)** 1,359 genes with functional *de novo* mutations from the healthy control dataset (**Supplementary Table 14**); **4)** 444 Well-known dominant ID genes (**Supplementary Table 4**).

### Gene set based evaluation of pLI

We evaluated the pLI by computing the expected median pLI for each gene set based on randomly drawing *n* pLI values from the complete set of 18,226 pLI annotated genes and calculate the median (where *n* is the number of genes in the gene set). By repeating this random sampling process 100.000 times, we can compute the likelihood of the observed median pLI to the expected median pLI by calculating the empirical p-value:

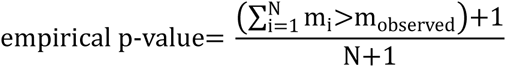

where *m* is the median pLI of one simulation, *m*_*observed*_ is the observed median pLI and *N* is the total number of performed simulations (*N*=100.000). Based on the simulations we identified a significant lower (Observed 9.33×10^−9^ vs. expected 0.03; empirical p-value: <1×10^−5^) median pLI for the loss-of-function tolerant (LoFT) genes which is in line with the LoF tolerant nature of this gene set. For the healthy control set the observed median pLI matched the expected median pLI (Observed 0.03 vs. expected 0.03; empirical p-value: 0.31). For the “house-keeping” and dominant ID gene sets the observed median pLI is significantly higher than the expected median pLI (Observed: 0.87 vs. expected: 0.03; empirical p-value =1×10^−5^; Observed: 0.95 vs. expected: 0.03; empirical p-value <1×10^−5^, respectively). The median pLI of the “house-keeping” gene approximates (median pLI = 0.87) and the dominant ID gene set (median pLI = 0.95) surpasses the extremely LoF intolerant threshold of 0.9 which is in line with the LoF intolerant nature for of “house-keeping” and dominant ID genes (**Supplementary Figure 12**).

The set of ten novel candidate ID genes has a median pLI of 0.99 (Observed 0.99 vs. expected 0.05; empirical p-value <1x10^−5^) which is, as observed for the dominant ID genes, above the extremely LoF gene threshold of 0.9 (**Supplementary Figure 8**). For the 21 dominant ‘missense only’ genes (with at least 3 missense mutations in the absence of LoF mutations) we observe the highest median pLI of 0.9999 (Empirical p-value <1×10^−5^) illustrating that those known and novel candidate dominant ID genes that harbor only missense variants are among the most LoF intolerant ID genes (**Supplementary Figure 8**).

### Attributing Residual Variation Intolerance Score (RVIS) for all genes

In addition, the Residual Variation Intolerance Score (RVIS) were assessed for the same gene sets as for the aforementioned pLI. The RVIS ranks genes based on whether they have more or less common functional genetic variation relative to the genome-wide expectation. The initial RVIS gene scores were computed based on the NHLBI-ESP6500 data set^20^ and recently recomputed based on the ExAC v0.3 dataset (http://genic-intolerance.org/). The genes from our study were annotated with the RVIS scores based on ExAC (**Supplementary Table 8**).

RVIS scores for gene sets were compared in the same way as for the pLI (**Supplementary Figure 13**). Again, we found the set of ten novel candidate ID genes to be significantly more intolerant than any random set of genes found (empirical p-value = 4.60×10^−4^), similar to the known dominant ID genes (**Supplementary Figure 13**). For the 21 dominant ‘missense only’ genes we again observe the lowest median RVIS of 3.56 (Empirical p-value <1×10^−5^; **Supplementary Figure 13**).

### Estimating clustering of *de novo* mutations

The spatial distribution of missense, frame-shifts and nonsense DNMs were analyzed for clustering within the respective gene they occurred based on 100,000 simulations. The locations of observed DNMs were randomly sampled over the coding exons of the gene and the distances (in base pairs) between the mutations were normalized for the total coding size of the respective gene. The geometric mean (the *n*^th^ root of the product of *n* numbers) of all mutation distances between the DNMs was taken as a measure of clustering. A pseudo count (adding one to all distances and one to the gene size) was applied to avoid a mean distance of 0 when there are identical mutations and ignoring the distances to the remaining DNMs.

Based on the prior distance distribution of the 100,000 simulations, a gene-based empirical probability of the observed distance was computed for dominant ID genes with 3 or more DNMs (n=64 genes) in the ID set of 2,104 trios. A total of 21 genes contained only missense mutations (“missense only” group) and 43 genes contained frame-shift, nonsense or a combination of frame-shift, nonsense and missense DNMs (“LoF + Functional” group). In 21 genes of the “missense only” group five genes had an empirical probability below the significant threshold of 0.05/64, whereas only one of the 43 “LoF + Functional” genes had a empirical probability below the significant threshold (**Supplementary Table 9**). Fisher’s exact test was used to compute the statistical significance.

### Clinical evaluation of selected patients

All patients were referred by clinical geneticists for diagnostic evaluation and overall patient characteristics were comparable to a previously published cohort^13^. To confirm the identification of the novel candidate ID genes, we compared the phenotypes of individuals with a DNM in any one of the 10 novel candidate genes and two genes (*SLC6A1 and TCF7L2*) significantly enriched in the neurodevelopmental cohort. Comparison of phenotypes was only possible for 8/12 genes in which at least two individuals with ID were in the RUMC cohort (7/10 novel candidate ID genes, and 1/2 novel candidate NDD genes). Detailed clinical information of other published individuals is mostly not available. A table listing these clinical details is provided in **Supplementary Table 7**. For *TLK2* and *SETD2* a more detailed phenotypic comparison was performed (see case reports below).

## Acknowledgments

The authors would like to thank the Exome Aggregation Consortium and the groups that provided exome variant data for comparison. A full list of contributing groups can be found at http://exac.broadinstitute.org/about. We thank all clinicians involved for referring individuals with ID for diagnostic exome sequencing. We thank Jelle Goeman for statistical advice and Matthew Hurles for useful discussions. We would also like to thank the participating individuals and their families. This work was in part financially supported by grants from the Netherlands Organization for Scientific Research (912-12-109 to J.A.V., A.S. and B.B.A.d.V., 916-14-043 to C.G., 907-00-365 to T.K. and SH-271-13 to C.G. and J.A.V.) and the European Research Council (ERC Starting grant DENOVO 281964 to J.A.V.).

## Supplementary Text

**Case reports of individuals with DNMs in selected novel candidate ID genes**

### TLK2

#### Patient 17

This male is the second of three children of consanguineous parents of Turkish ancestry. His father had learning difficulties and was illiterate. He was born after 40+3 weeks of gestation with a birth weight of 3415 gram (0 SD), length of 50 cm (-2 SD) and head circumference of 35 cm (0 SD). Delivery was uncomplicated. Both motor and language development were delayed. He started walking at the age of 2 years. At the age of 20 years, psychological assessment revealed a TIQ level of 56 (WISC-III). He followed special education as a child, but after finishing this, he was not able to work due to severe psychiatric problems with frequent tantrums and periods of hyperventilation. Several psychiatric medications turned out to be insufficient. Ophthalmologic examination revealed myopia (-2.5/-3.5 dioptre) and strabismus of the left eye. He was treated for oesophagitis and had frequent diarrhea. Because of his delayed development, he was referred to a clinical geneticist. Physical examination at the age of 16 years showed a normal height of 171 cm (-0.5 SD, based on normal values for Turkish children^1^ and normal weight of 50.3 kg (-0.5 SD). He had facial dysmorphisms including short forehead, upward slanting palpebral fissures, mild epicanthal folds, hypertelorism, ptosis, flat mid-face, thin upper lip and pointed chin. Thoracal kyphosis and scoliosis were noticed. His hands showed contractures of the proximal interphalangeal joints of the fourth fingers and absence of flexion creases of the distal interphalangeal joints of the second till fourth fingers. Previous investigations, comprising of karyotyping, fragile X screening and SNP array analysis were normal. Metabolic screening and brain MRI scan, performed at age 16 years, showed no abnormalities. Using whole exome sequencing, a *de novo TLK2* mutation was identified: Chr17(GRCh37):g.60678182G>T; NM_006852.3:c.1720+1G>T (p.(r.spl?)).

#### Patient 439

This male patient is the second of two children of nonconsanguineous parents of Dutch ancestry. There were no developmental problems reported in the family. Prenatal ultrasound showed a single umbilical artery. He was born after 40 weeks of gestation with a birth weight of 3170 gram (-1 SD) after an uncomplicated delivery. As neonate, he had feeding difficulties and low weight. His development was delayed. He started walking at the age of 18 months. He spoke the first words at normal age, but further language development was delayed and he speaks with hoarse voice. Psychological assessment at the age of 5 years showed a borderline TIQ level of 80 and at the age of 7.5 years a mild intellectual disability with a TIQ level of 67 (SON-R 6-40). He was recently diagnosed with multiple complex developmental disorders, with severe problems in regulation of emotions and anxiety. Treatment with Abilify and Cipramil has been started to alleviate the symptoms. The boy had severe constipation, requiring laxatives. Physical examination at the age of 7 years showed a short stature, with a height of 11.5 cm (-2.5 SD), normal weight of 21 kg (0 SD) and normal head circumference of 50 cm (-1 SD). He had facial dysmorphisms including upward slanting palpebral fissures, blepharophimosis, ptosis, full nasal tip and pointed chin. His hands showed brachydactyly and single flexion crease of the second and third finger. Previous investigations, consisting of fragile X screening and SNP array, were normal. Metabolic screening showed no abnormalities. Using whole exome sequencing a *de novo TLK2* mutation was identified: Chr17(GRCh37):g.60689765C>T; NM_006852.3:c.2092C>T (p.(Arg698*)).

### SETD2

#### Patient 162

This male patient is the first child of three children of non-consanguineous parents of Dutch ancestry. There were no developmental problems reported in the family. He was born after 36+4 weeks of gestation which was complicated by maternal hypertension and HELLP syndrome. Delivery was uncomplicated. His birth weight of 2625 gram (-1.5 SD) was normal. There was normal motor development, but delayed development of language, with the first words at the age of 3 year. At the age of 7 years, psychological assessment showed mild ID with TIQ of 54 (WISC-III). As child, he received surgery for bilateral inguinal hernia. He needed glasses to correct myopia (-2/-2.5 dioptre). Physical examination at the age of 11 years showed macrocephaly with a head circumference of 59.5 cm (+4 SD), high-normal height of 164.4 cm (+2 SD) and low-normal weight of 45 kg (-2 SD). He had facial dysmorphisms including small and protruding ears, broad forehead, hypertelorism, downward slanting and narrow palpebral fissures. A shawl scrotum was present. There were no abnormalities of the extremities observed. At age of 4 years, brain MRI and metabolic screening showed no abnormalities. Previous investigations, consisting of karyotyping, MLPA analysis, fragile X screening, 250k SNP array and sequence analysis of multiple genes (*SOS1, NRAS, SHOC2, NF1, SPRED1, NSD1, KCNQ1OT1, H19, NFIX, EZH2, BRAF, MAP2K1, KRAS, PTPN11, RAF1*) were normal. Using whole exome sequencing a *de novo SETD2* mutation was identified: Chr3(GRCh37):g.47161721dup; NM_014159.6:c.4405dup (p.(Met1469fs)).

#### Patient 716

This male patient was the second of two children of non-consanguineous parents of Dutch ancestry. His older brother needed special education because of severe behavior problems. Further family history was not contributory. He was born after 40 weeks of gestation with a birth weight of 3000 gram (-1.5 SD). Delivery was uncomplicated. In the first year, no problems were noticed, but after his first year, both motor and language development were delayed. He walked the first steps at the age of 19 months. As young child, he was aggressive to other children, but this improved from the age of 8 years. He was referred to a clinical geneticist because of developmental delay, tall stature and macrocephaly. From age 9 years, he eats unrestrained and gains weight. Physical examination at the age of 9 years showed tall stature with height of 162.5 cm (+2.5 SD), macrocephaly with head circumference of 57.8 cm (+3 SD) and normal weight of 55.5 kg (+1.5 SD). He had facial dysmorphisms including high forehead, deep set eyes and downward slanting of palpebral fissures. There was clinodactyly of his second and third toes and he had a mastocytoma on his right leg. Brain MRI at the age of 9 years showed a subarachnoid cyst left temporal. Analysis of growth hormones showed no abnormalities, but hand films revealed advanced carpal and phalangeal bone age of 11 years at the age of 9 years. Previous investigations, consisting of karyotyping, fragile X and Sotos screening were normal. Using family-based whole exome sequencing a *de novo* mutation in *SETD2* was identified: Chr3(GRCh37):g.47155435_47155437del; NM_014159.6:c.4644_4646del (p.(Gln1548del)).

**Supplementary Tables**

**Supplementary Table 1.** **Meta-study cohort composition**.

| Study | # Trios | Disorder | Coding | Functional | LoF | Ref |
| --- | --- | --- | --- | --- | --- | --- |
| RUMC | 820 | ID | 1,083 | 872 | 211 |  |
| Gilissen *et al.* 2014 | 50 | ID | 84 | 65 | 15 | <sup>2</sup> |
| Rauch *et al.* 2012 | 51 | ID | 84 | 76 | 21 | <sup>3</sup> |
| de Ligt *et al.* 2012 | 50* | ID | 57 | 46 | 12 | <sup>4</sup> |
| DDD. 2014 | 1,133 | ID | 1,337 | 1,073 | 235 | <sup>5</sup> |
| Neale *et al.* 2012 | 175 | ASD | 170 | 120 | 18 | <sup>6</sup> |
| Iossifov *et al.* 2014 | 2,508 | ASD | 2,738 | 2,103 | 389 | <sup>7</sup> |
| Epi4K. 2014 | 356 | EE | 412 | 333 | 53 | <sup>8</sup> |
| Xu *et al.* 2011 | 53 | SCZ | 34 | 32 | 0 | <sup>9</sup> |
| McCarthy *et al.* 2014 | 57 | SCZ | 64 | 50 | 9 | <sup>10</sup> |
| Gulsuner *et al.* 2013 | 105 | SCZ | 99 | 67 | 12 | <sup>11</sup> |
| Xu *et al.* 2012 | 231 | SCZ | 132 | 106 | 19 | <sup>12</sup> |
| Fromer *et al.* 2014 | 617 | SCZ | 639 | 483 | 63 | <sup>13</sup> |
| **Total** | **6,206** |  | **6,933** | **5,426** | **1,057** |  |

Columns indicate (from left to right) the study name, the number of trios in the studied cohort, the disorder that was studied (ID: Intellectual disability, SCZ: Schizophrenia ASD: autism spectrum disorder, EE: epileptic encephalopathy), the number of coding, functional and loss-of-function (LoF) mutation. *Of the 100 samples in de Ligt *et al.* 50 samples overlap with Gilissen *et al.* Overlapping DNMs are removed from the de Ligt *et al.* DNM list.

**Supplementary Table 2. All identified *de novo* mutations in the RUMC cohort**

(Excel file)

**Supplementary Table 3. Genes significantly enriched for *de novo* mutations in the full RUMC cohort**

(Excel file)

**Supplementary Table 4. Lists of known ID genes**

(Excel file)

**Supplementary Table 5.Gene stats for the RUMC set**

(Excel file)

**Supplementary Table 6.Gene stats for the ID set**

(Excel file)

**Supplementary Table 7.** **Phenotypes of patients with a mutation in a novel candidate ID gene**.

|  |  |  |  |  |  |  |  |  |  |  |  |  |  |  |  |  |  |  |  |
| --- | --- | --- | --- | --- | --- | --- | --- | --- | --- | --- | --- | --- | --- | --- | --- | --- | --- | --- | --- |
| TCF20 | 530 | + (Mild) |  |  |  |  |  |  | + |  |  |  |  |  |  | < -2.5 SD |  |  | > 2.5 SD |
|  | 336 | + (Moderate) |  | + | + | + |  |  | + |  |  |  |  |  |  |  |  |  |  |
|  | 720 | + (Moderate) |  |  | + |  | + |  |  |  |  |  |  |  |  |  | < -2.5 SD | < -2.5 SD | < -2.5 SD |
|  | 751 | + (Mild) |  |  | + |  |  |  | + |  |  |  | + |  |  |  |  |  | < -2.5 SD |
| TLK2 | 439 | + (Mild) |  | + |  |  | + |  | + | + |  |  | + |  |  |  |  | < -2.5 SD |  |
|  | 17 | + (Mild) |  | + | + |  | + |  | + | + | + |  | + |  |  |  |  |  |  |
| TRIP12 | 188 | + |  |  | + | + |  |  | + |  |  |  |  |  | + | + |  |  |  |
|  | 24 | + (Moderate) |  |  |  |  |  |  |  |  |  |  |  |  |  |  | > 2.5 SD |  |  |

**Supplementary Table 8. List of used genes and corresponding pLI and RVIS**

(Excel file)

**Supplementary Table 9. Clustering of mutations in genes with only missense mutations**

(Excel file)

**Supplementary Table 10.Gene stats for the NDD set**

(Excel file)

**Supplementary Table 11.**
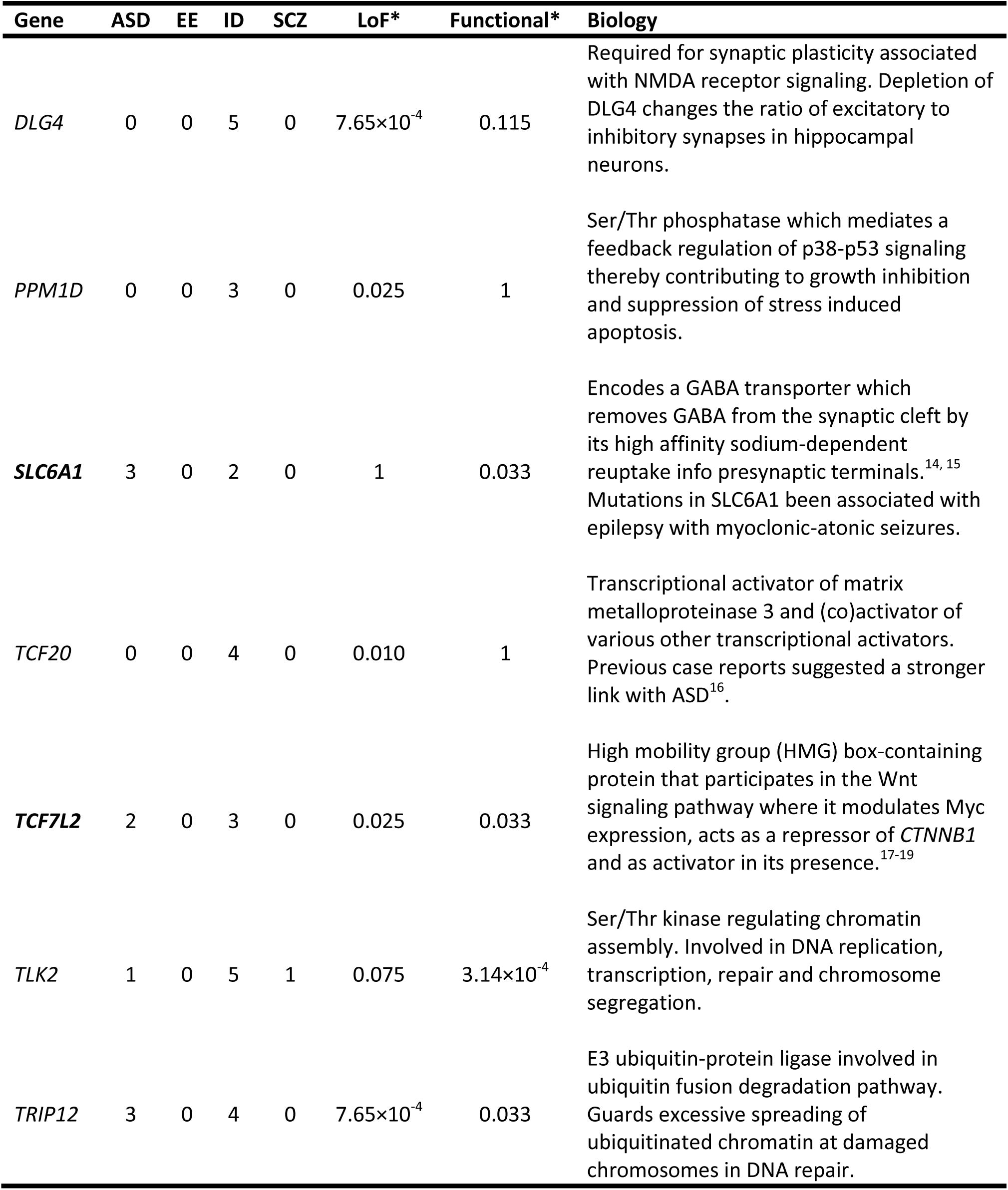
**All significant genes in the neurodevelopmental cohort**.

| Gene | ASD | EE | ID | SCZ | LoF* | Functional* | Biology |
| --- | --- | --- | --- | --- | --- | --- | --- |
| *DLG4* | 0 | 0 | 5 | 0 | $7.65 \times 10^{-4}$ | 0.115 | Required for synaptic plasticity associated with NMDA receptor signaling. Depletion of DLG4 changes the ratio of excitatory to inhibitory synapses in hippocampal neurons. |
| *PPM1D* | 0 | 0 | 3 | 0 | 0.025 | 1 | Ser/Thr phosphatase which mediates a feedback regulation of p38-p53 signaling thereby contributing to growth inhibition and suppression of stress induced apoptosis. |
| *SLC6A1* | 3 | 0 | 2 | 0 | 1 | 0.033 | Encodes a GABA transporter which removes GABA from the synaptic cleft by its high affinity sodium-dependent reuptake into presynaptic terminals.<sup>14, 15</sup> Mutations in SLC6A1 been associated with epilepsy with myoclonic-atonic seizures. |
| *TCF20* | 0 | 0 | 4 | 0 | 0.010 | 1 | Transcriptional activator of matrix metalloproteinase 3 and (co)activator of various other transcriptional activators. Previous case reports suggested a stronger link with ASD<sup>16</sup>. |
| *TCF7L2* | 2 | 0 | 3 | 0 | 0.025 | 0.033 | High mobility group (HMG) box-containing protein that participates in the Wnt signaling pathway where it modulates Myc expression, acts as a repressor of *CTNNB1* and as activator in its presence.<sup>17-19</sup> |
| *TLK2* | 1 | 0 | 5 | 1 | 0.075 | $3.14 \times 10^{-4}$ | Ser/Thr kinase regulating chromatin assembly. Involved in DNA replication, transcription, repair and chromosome segregation. |
| *TRIP12* | 3 | 0 | 4 | 0 | $7.65 \times 10^{-4}$ | 0.033 | E3 ubiquitin-protein ligase involved in ubiquitin fusion degradation pathway. Guards excessive spreading of ubiquitinated chromatin at damaged chromosomes in DNA repair. |

Genes with a significant enrichment for *de novo* mutations identified in the complete cohort of neurodevelopmental trios. Genes in bold were not identified in the ID cohort analysis. Columns show, from left to right, the gene name, number of cases found in Autism Spectrum Disorder (ASD) cohorts, in Epileptic Encephalopathy (EE) cohorts, in intellectual disability (ID) and Schizophrenia (SCZ). *Corrected p-value for loss-of-function (LoF) and functional mutations, respectively.

**Supplementary Table 12.** Control cohort composition. For each of the cohorts used to construct the control set the number of samples and *de novo* mutations is shown. Columns indicate (from left to right) the study name, the number of trios in the studied cohort, the disease that was studied in the context of the control cohort (ID: Intellectual disability, SCZ: Schizophrenia ASD: autism spectrum disorder, EE: epileptic encephalopathy) as well as whether samples were unrelated healthy controls (Con.) or unaffected siblings (Sibs.), the number of coding, functional and loss-of-function (LoF) mutation, the sequencing platform used, the study of Gulsuner *et al.* used the Illumina Genome Analyzer IIX where the other four studies used the Illumina HiSeq2000 platform, the enrichment, enrichment kits used, coverage, reported coverage as stated in the published paper.

| Study | # Trios | Disorder | Coding | Functional | LoF | Sequencer | Enrichment Kit | Coverage |
| --- | --- | --- | --- | --- | --- | --- | --- | --- |
| Iossifov *et al.* 2014<sup>(7)</sup> | 1,911 | ASD / Sibs. | 1,780 | 1,309 | 175 | CSHL: Hiseq2000 | SeqCap EZ v2.0(44 Mb) | 80% enriched targets at 20x coverage |
|  |  |  |  |  |  | UW: Hiseq2000 | SeqCap EZ v2.0(44 Mb) |  |
|  |  |  |  |  |  | YALE: HiSeq2000 | SeqCap EZ v2.0(44 Mb) |  |
| GoNL. 2014<sup>(20)</sup> | 250 | Con. | 137 | 97 | 7 | HiSeq2000 | none (WGS) | Average 13x |
| Rauch *et al.* 2012<sup>(3)</sup> | 20 | ID / Con. | 19 | 12 | 2 | HiSeq2000 | SureSelect XT Human all exon 50 Mb kit | Median coverage: 112x. 90% enriched targets at 20x |
| Gulsuner *et al.* 2013<sup>(11)</sup> | 84 | ID / Sibs. | 66 | 48 | 11 | Genome Analyzer IIX | SeqCap EZ v2.0 (44 Mb) | Median coverage >100x. 93% enriched targets at 10 x |
| Xu *et al.* 2012 <sup>(12)</sup> | 34 | SCZ / Con. | 17 | 12 | 1 | HiSeq2000 | SureSelect 37Mb | Median coverage:<br>65.2x. >80%<br>enriched targets<br>at 20x |
| **Total** | **2,299** |  | **2,019** | **1,478** | **196** |  |  |  |

**Supplementary Table 13.**
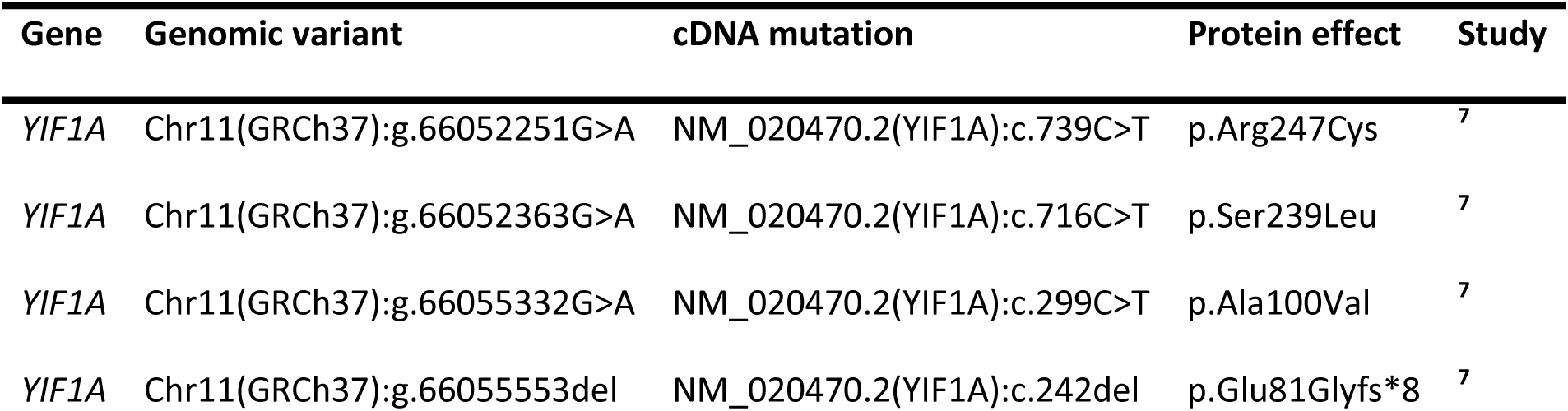
**List of DNMs in genes significantly enriched in the control cohort**.

| Gene | Genomic variant | cDNA mutation | Protein effect | Study |
| --- | --- | --- | --- | --- |
| *YIF1A* | Chr11(GRCh37):g.66052251G>A | NM_020470.2(YIF1A):c.739C>T | p.Arg247Cys | <sup>7</sup> |
| *YIF1A* | Chr11(GRCh37):g.66052363G>A | NM_020470.2(YIF1A):c.716C>T | p.Ser239Leu | <sup>7</sup> |
| *YIF1A* | Chr11(GRCh37):g.66055332G>A | NM_020470.2(YIF1A):c.299C>T | p.Ala100Val | <sup>7</sup> |
| *YIF1A* | Chr11(GRCh37):g.66055553del | NM_020470.2(YIF1A):c.242del | p.Glu81Glyfs*8 | <sup>7</sup> |

**Supplementary Table 14.Gene stats for the control set**

(Excel file)

**Supplementary Figure 1.**
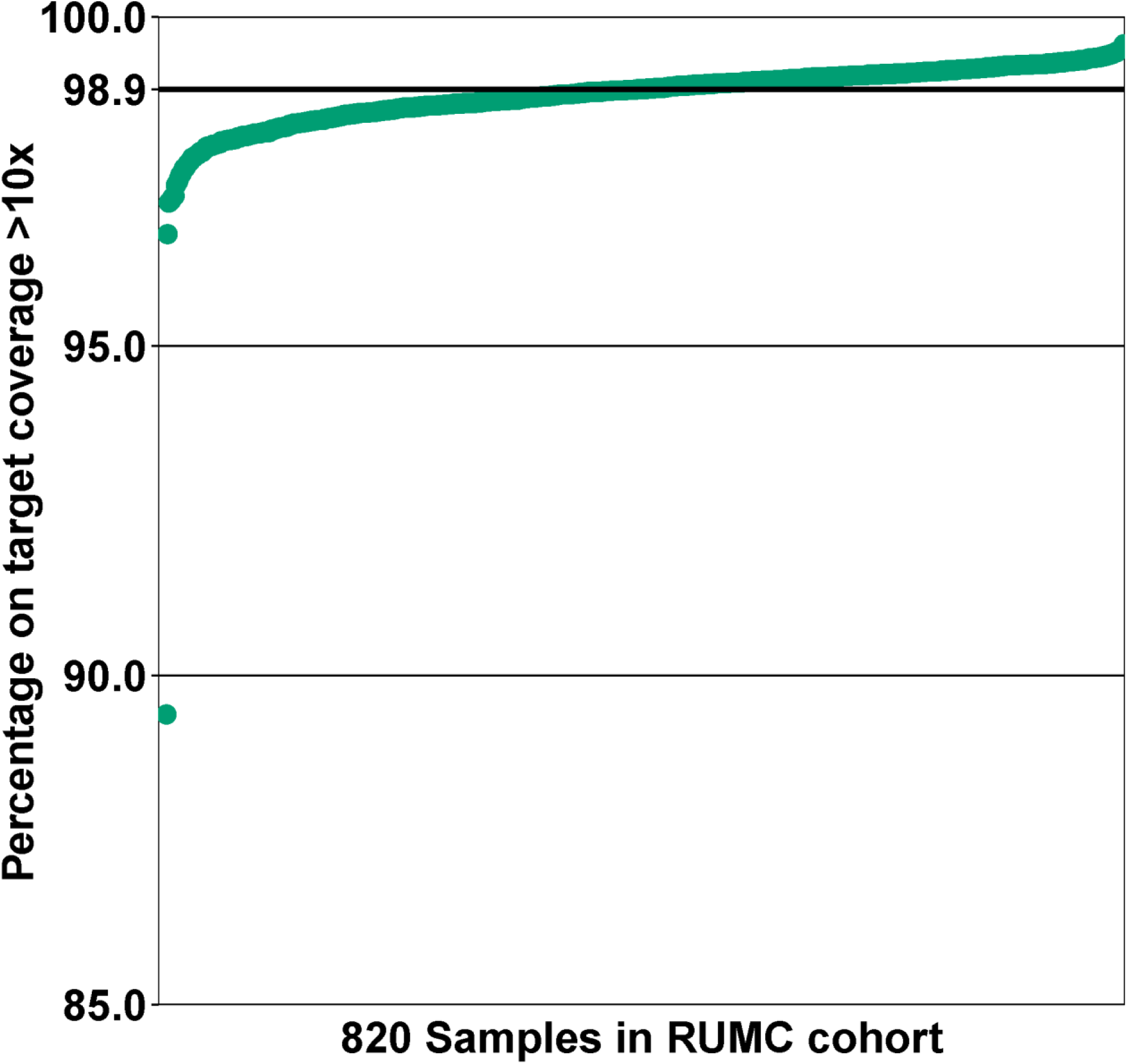
Overview of the percentage of on target coverage by 10 or more reads. A scatter plot of the percentage of on target coverage by 10 or more reads shown for the 820 samples of the RUMC cohort. In this figure the samples are ordered ascending based on the percentage of on target coverage. The median percentage of 98.9% on target coverage by 10 or more reads of the RUMC cohort is depicted by the black thick line.

**Supplementary Figure 2.**
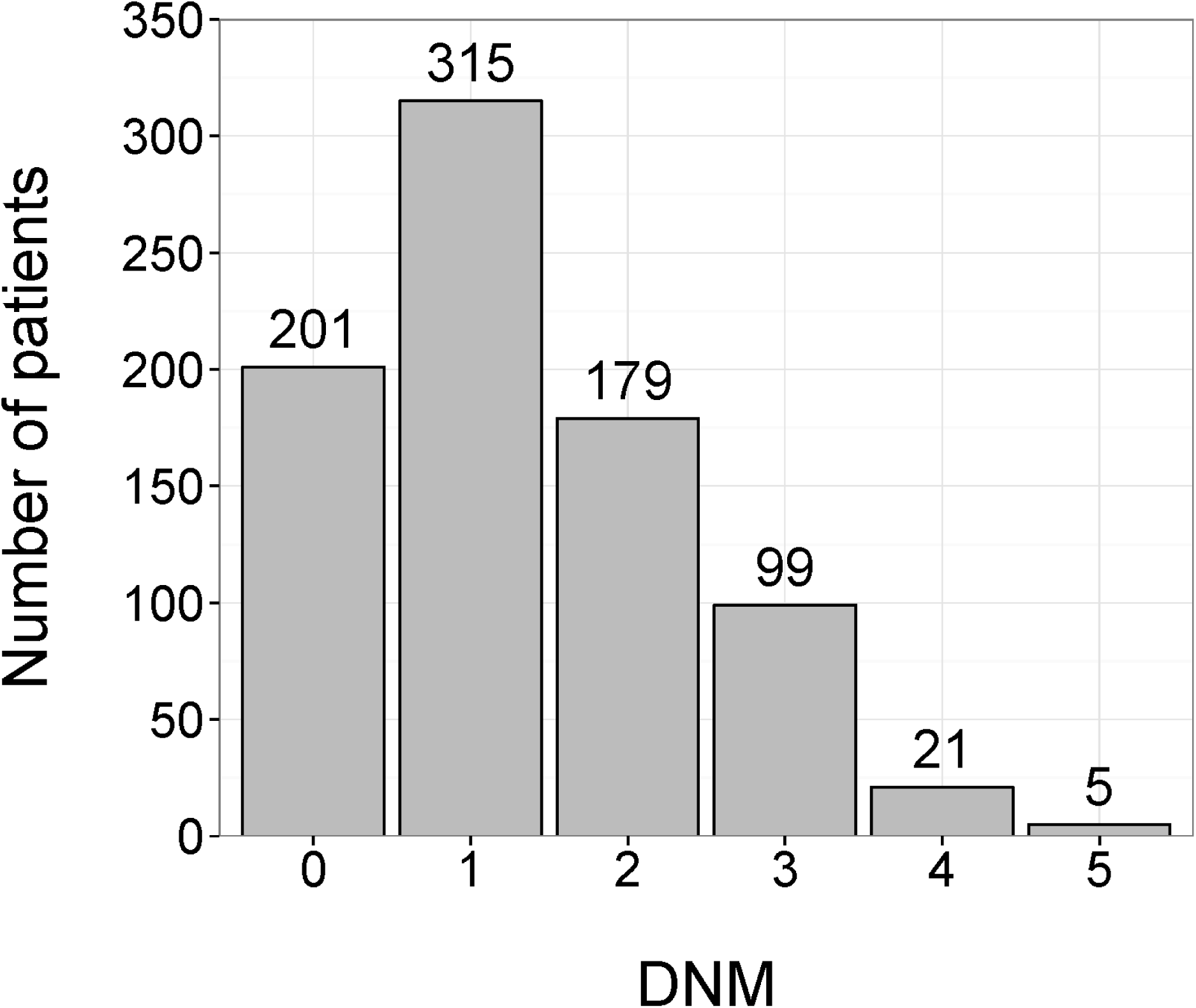
Distribution of *de novo* mutations (DNM) over patients in the RUMC cohort. In total, 619/820 patients had at least one *de novo* mutation.

**Supplementary Figure 3.**
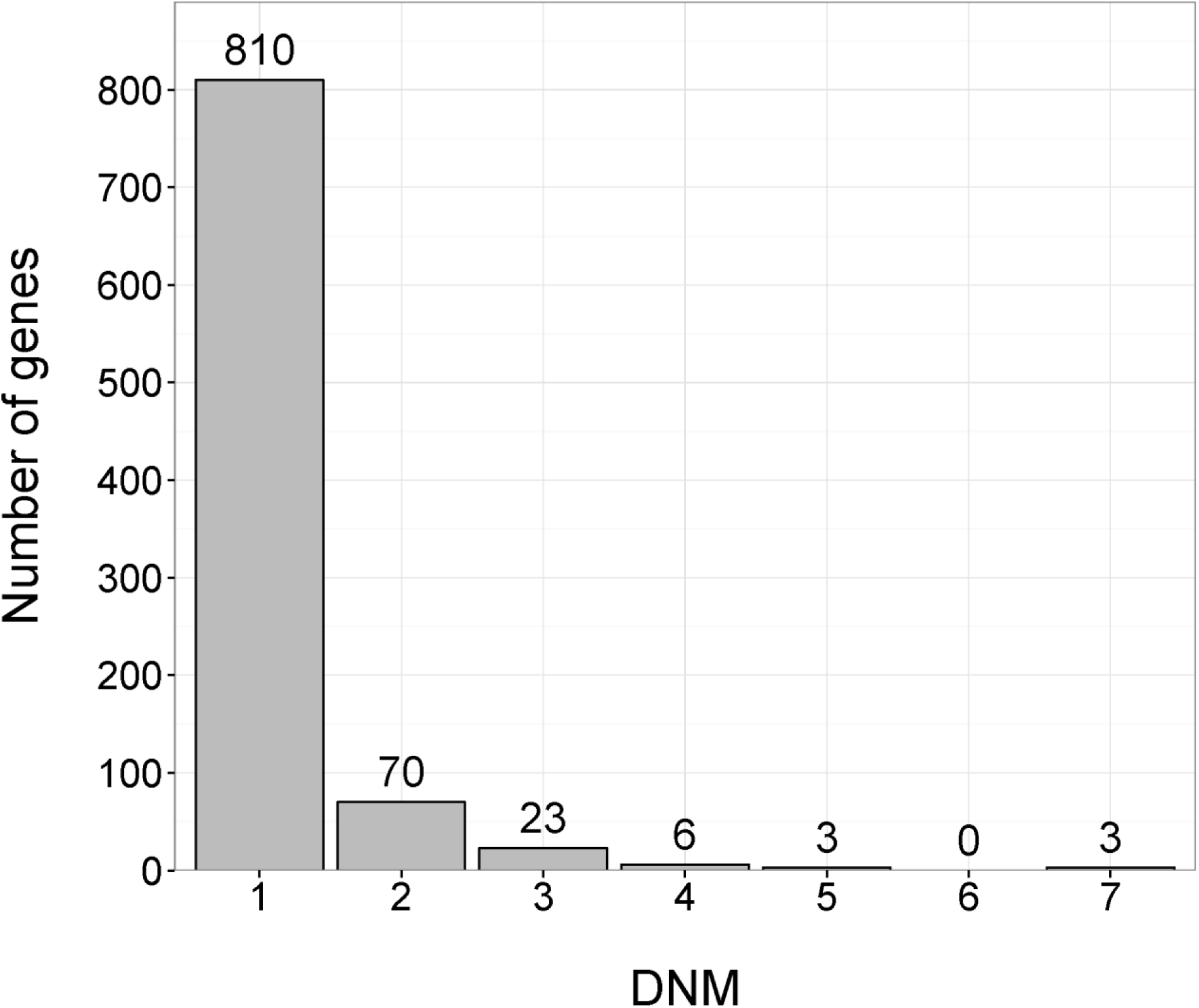
Distribution of *de novo* mutations per gene in the RUMC cohort. In total, 915 different genes have had at least one *de novo* mutation.

**Supplementary Figure 4.**
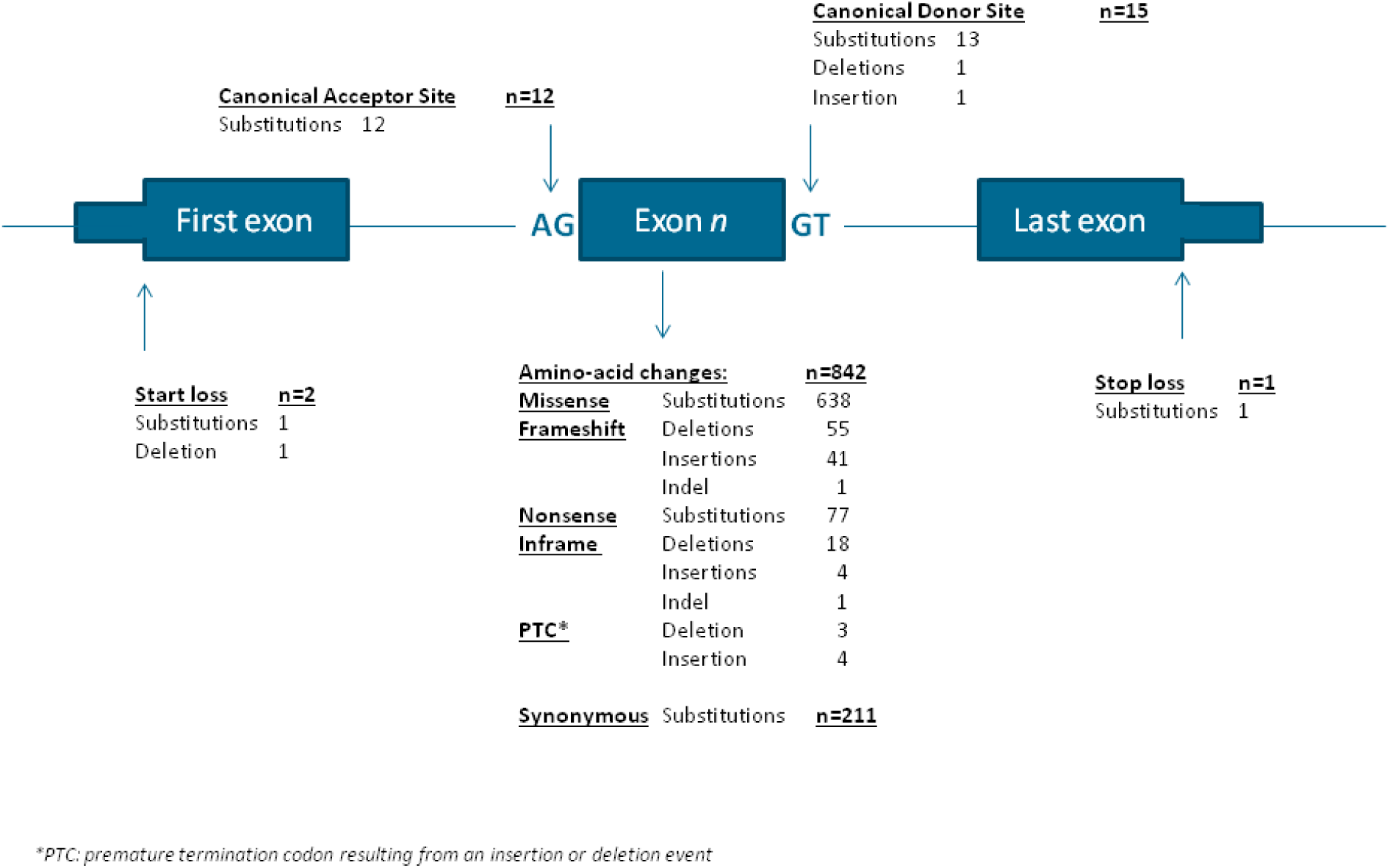
Schematic representation of the location of *de novo* mutations identified in the RUMC cohort and their presumed effect on protein function. *: Premature Termination Codon (PTC); An insertion or deletion does not introduce a frameshift event, but directly creates a PTC.

**Supplementary Figure 5.**
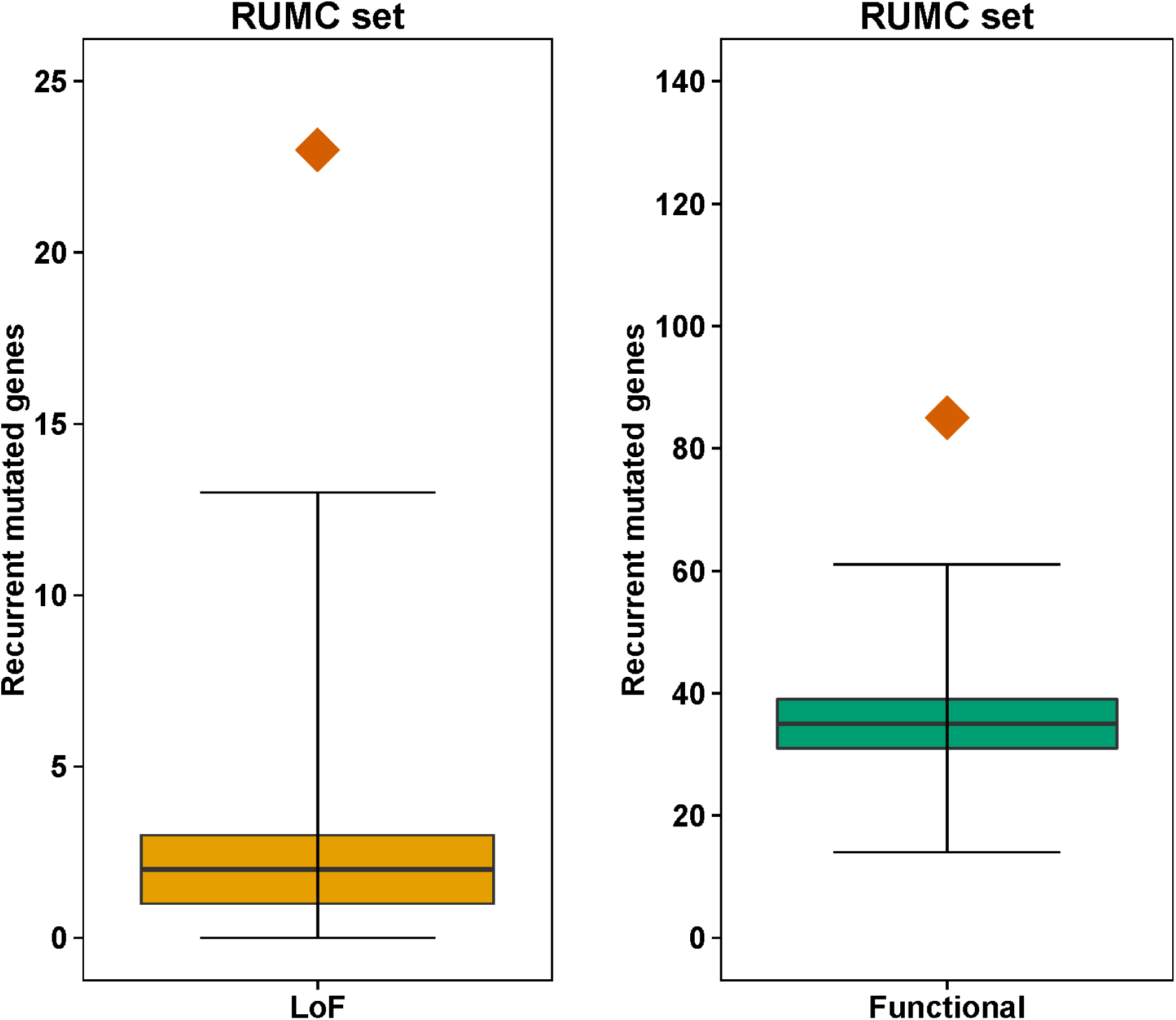
Simulations of recurrent mutated genes of the RUMC cohort. The panels show the distribution of the number of recurrently mutated genes based on 100,000 simulations. The colored boxes indicate the interquartile range; the whiskers indicate the full interval (LoF median: 2, range [0,13]; functional median: 35, range [14,61]) and the orange diamond indicate the observed number of recurrent *de novo* mutated genes in 820 ID patients (RUMC-cohort). For the LoF and the functional categories the observed number of recurrently mutated genes (observed LoF: 23; functional: 85) do statistically differ from the simulations based on the gene specific mutation rates of Samocha *et al.* (LoF empirical P-value: <1.00×10^−05^; functional empirical P-value: <1.00×50^−05^).

**Supplementary Figure 6.**
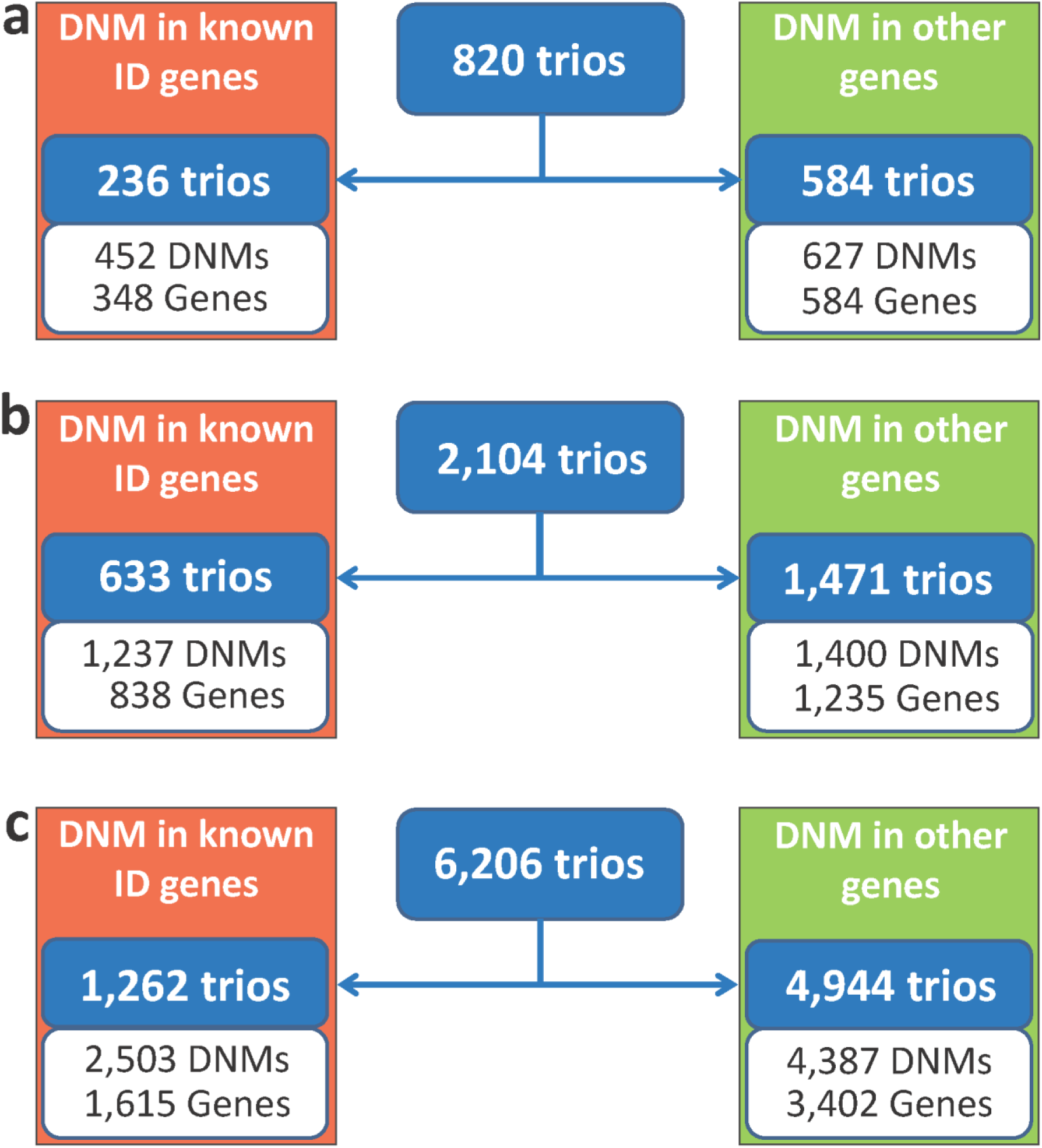
Flow chart of cohort construction for statistical analysis. Based on the presence of de novo mutations (DNMs) in 1,537 known ID genes (**Supplementary table 4**) the patients were divided among two groups for **a)** the RUMC cohort (820 trios) **b)** the combined ID cohort (2,104 trios) and **c)** the neurodevelopmental cohort (6,206 trios). The group on the left side in the color red indicate the patients with DNMs found in known ID genes. On the right, in color green, is the group of patients without DNMs present in the 1,537 known ID genes. The statistical analysis was performed on the cohort consisting of patients without DNMs in known genes. The number of trios and DNMs present in genes is shown for each group.

**Supplementary Figure 7.**
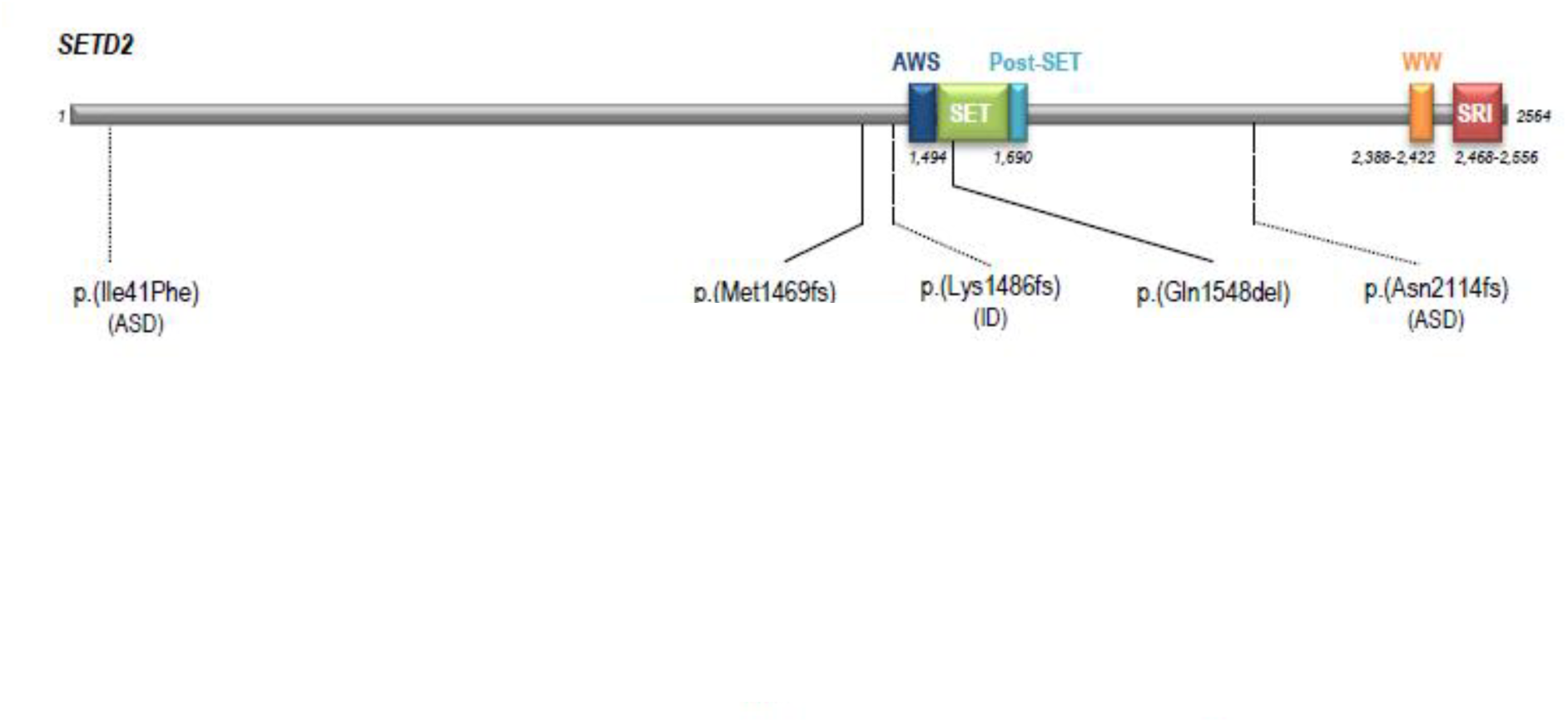
SETD2 (Q9BYW2) with *de novo* mutations in individuals with ID and ASD. Individuals show similar overgrowth phenotype including macrocephaly, tall stature and facial dysmorphisms **(Supplemental Case reports)**.

**Supplementary Figure 8.**
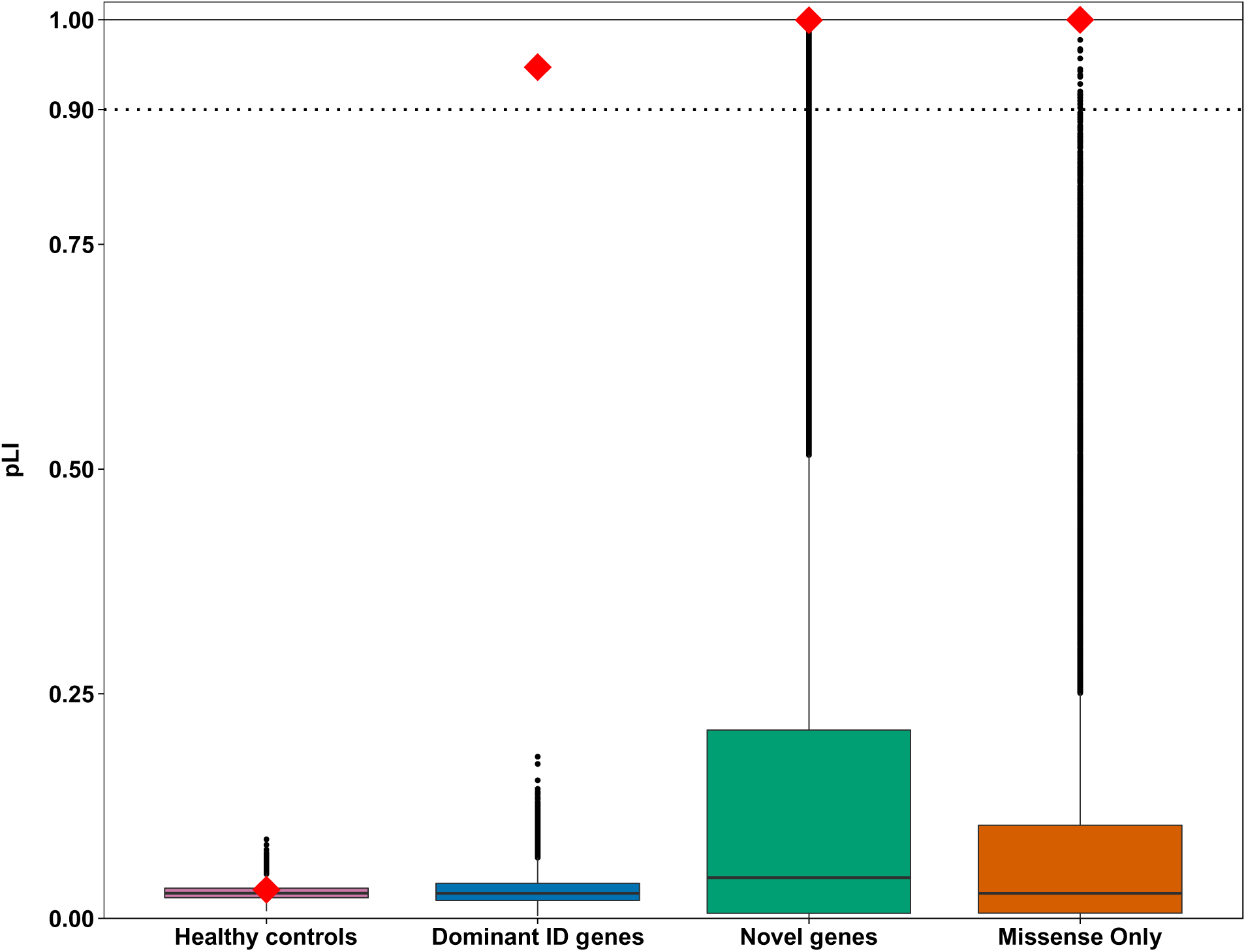
Intolerance to Loss-of-function (LoF) variation for ID genes. The box plots indicate the median pLI (probability LoF intolerant) for the gene sets based on 100,000 simulation (see **Online Methods**). The red diamonds show the observed median pLI. The closer a pLI is to 1 the more intolerant a gene is to LoF variants. A pLI >= 0.9 is considered as an extremely LoF intolerant set of genes^21^. The genes with at least one functional *de novo* mutation in healthy control set (Pink box; N=1,359) the observed median pLI matched the expected median pLI (Observed 0.03 vs. expected 0.03; empirical p-value: 0.31). For the set of dominant ID genes (blue box; N=9444) and ten novel candidate ID genes (green box) the observed median pLI is significantly higher than the expected median pLI (Observed: 0.87 vs. expected: 0.03; empirical p-value <1×10^−5^; Observed 0.99 vs. expected 0.05, respectively). For the dominant ‘missense only’ genes (with at least 3 missense mutations in the absence of LoF mutations; Orange box; N=21) we observe the highest median pLI of 0.9999 of all evaluated gene sets.

**Supplementary Figure 9.**
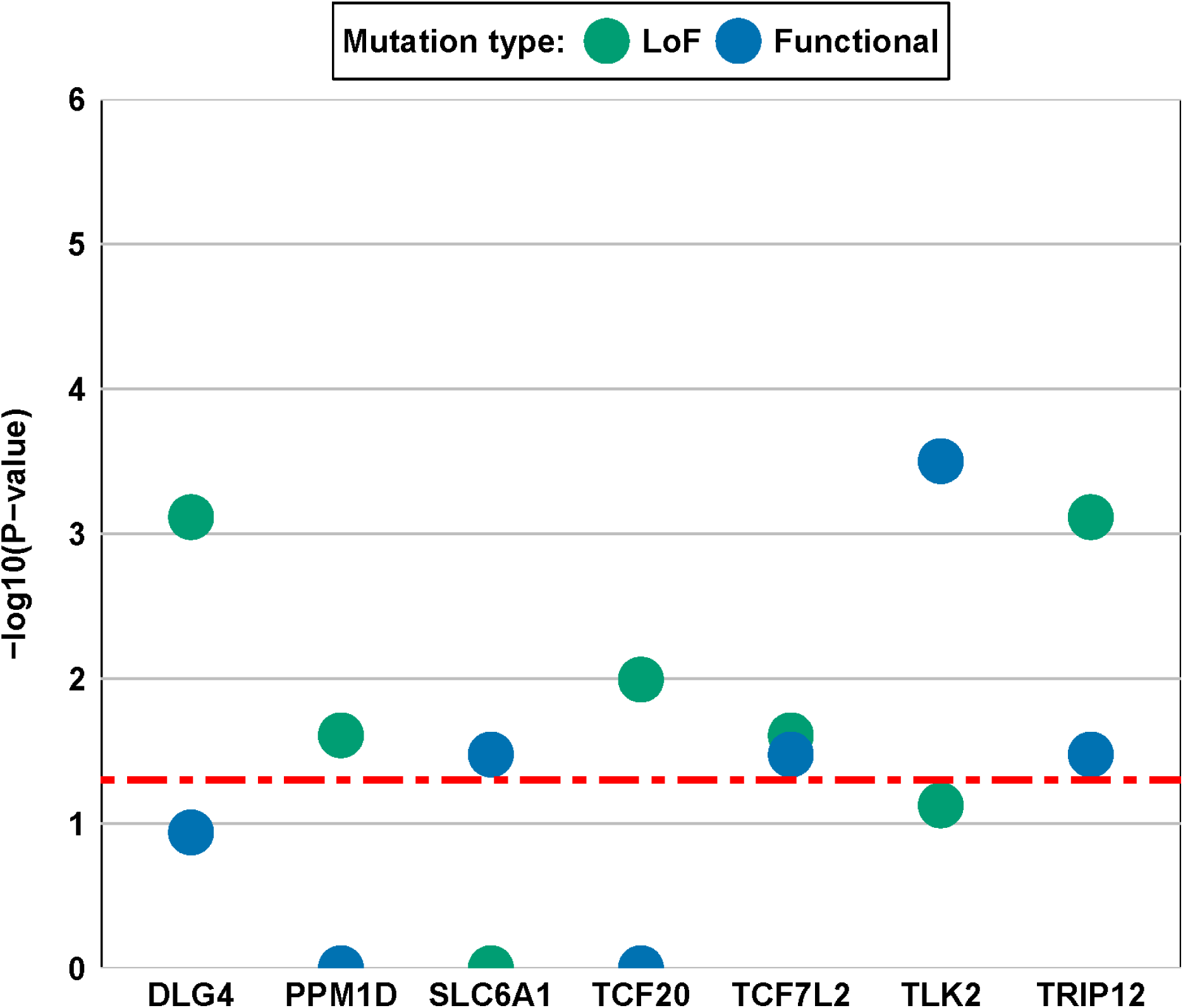
Genes enriched for LoF and functional *de novo* mutations in the cohort of 6,206 individuals with neurodevelopmental disease. The y-axes shows the-log10(P) value of the mutation enrichment. Corrected P-values based on LoF mutations are colored in green and corrected P-values based on functional mutations are colored blue. Only genes with a corrected P-value (LoF, functional, or both) less than the significance threshold (red dotted line, 0.05) are shown.

**Supplementary Figure 10.**
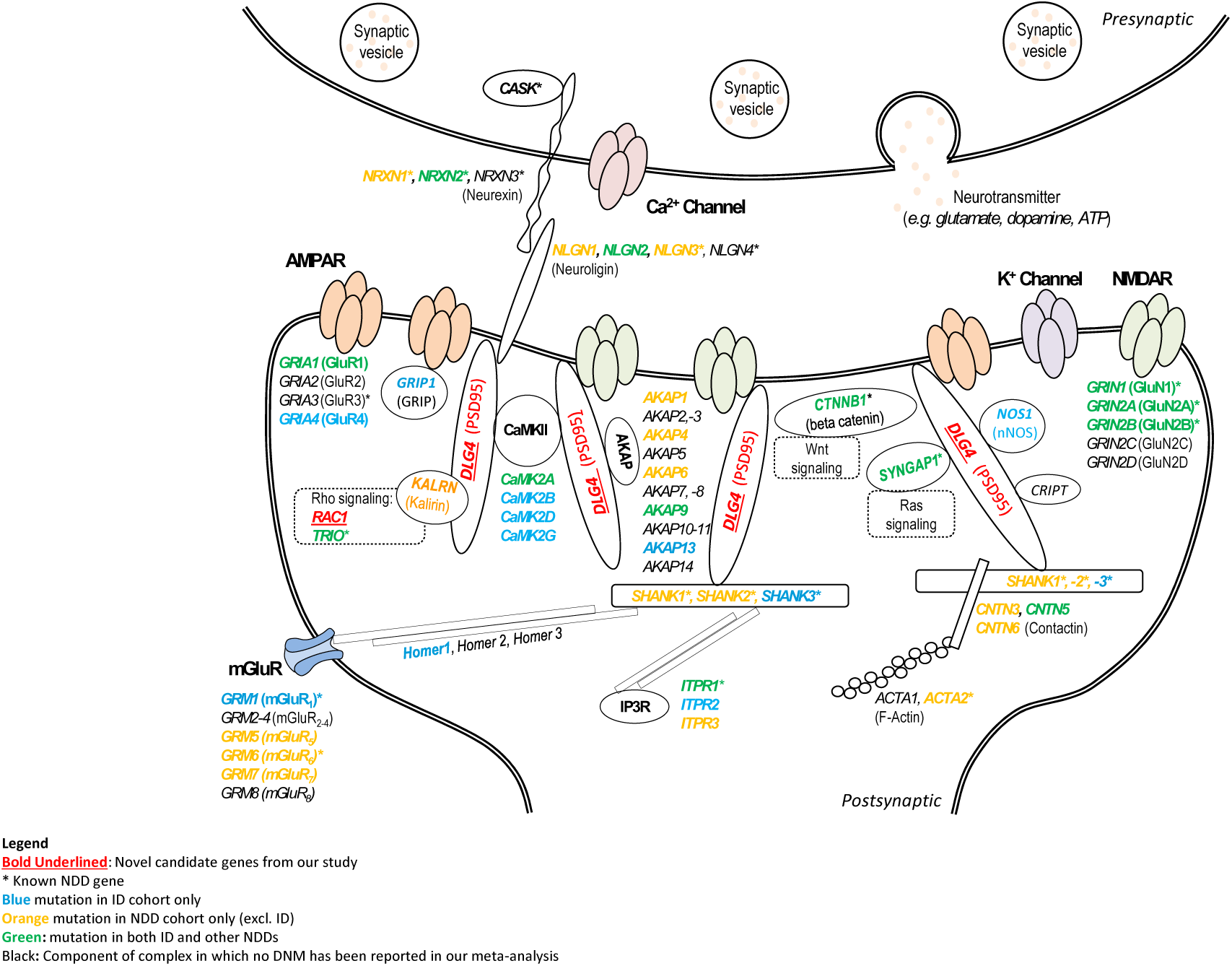
Schematic representation of a synapse, with special focus on the postsynaptic density (PSD). Proteins playing an essential role in the PSD for signaling cascades and/or receptor trafficking are schematically depicted, as well as the AMPA Receptor (AMPAR), NMDA receptor (NMDAR), metabolic Glutamate receptor (mGluR), Calcium and Potassium channels (Ca^2+^ and K^+^ respectively)^22^. An overlay was made between all DNMs identified in the NDD meta-analysis and the genes playing essential roles in the PSD, as well as with the list of known ID genes. Known ID genes are indicated by an asterisk. For genes in blue, we identified at least one DNM in the ID cohort, whereas the genes in orange were restricted to carry DNMs in the EE, SCZ and/or ASD patients. For genes in green, we identified DNMs in our meta-analysis both the ID and (at least one of the) NDD cohorts. Genes listed in black play a role in e.g. complex formation of the AMPAR or NMDAR, but in have not been identified to carry DNM in our current ID/NDD cohort. Importantly, three of ten genes which we identified as novel candidate ID gene play a role in the PSD and its downstream processes. *DLG4*, encoding post-synaptic density protein 95 (PSD95), is one of the core PSD proteins^23^, whereas *RAC1* and *TCF7L2* are important in downstream signaling cascades, including Rho-and Wnt signaling respectively (novel candidate ID are underlined and highlighted in red).

**Supplementary Figure 11.**
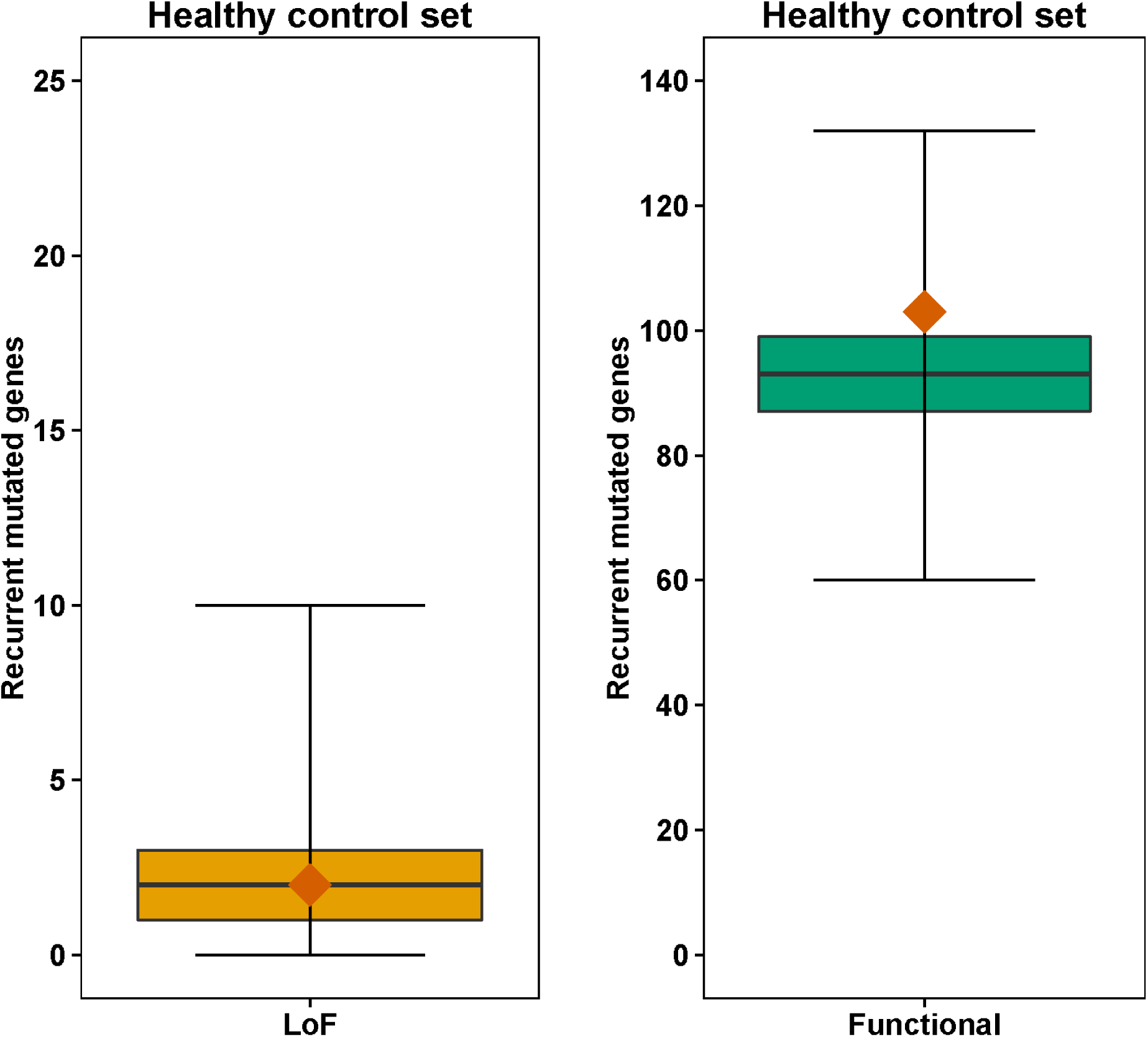
Simulations of recurrent mutated genes of the control cohort. The panels show the distribution of the number of recurrently mutated genes based on 100,000 simulations. The colored boxes indicate the interquartile range; the whiskers indicate the full interval (LoF median: 2, range [0,10]; functional median: 93, range [60,132]) and the orange diamond indicate the observed number of recurrent *de novo* mutated genes in 2,299 healthy controls. For the LoF and the functional categories the observed number of recurrently mutated genes (observed LoF: 2; functional: 103) do not statistically differ from simulations based on the gene specific mutation rates of Samocha *et al.* (LoF empirical P-value: 0.60; functional empirical P-value: 0.13).

**Supplementary Figure 12.**
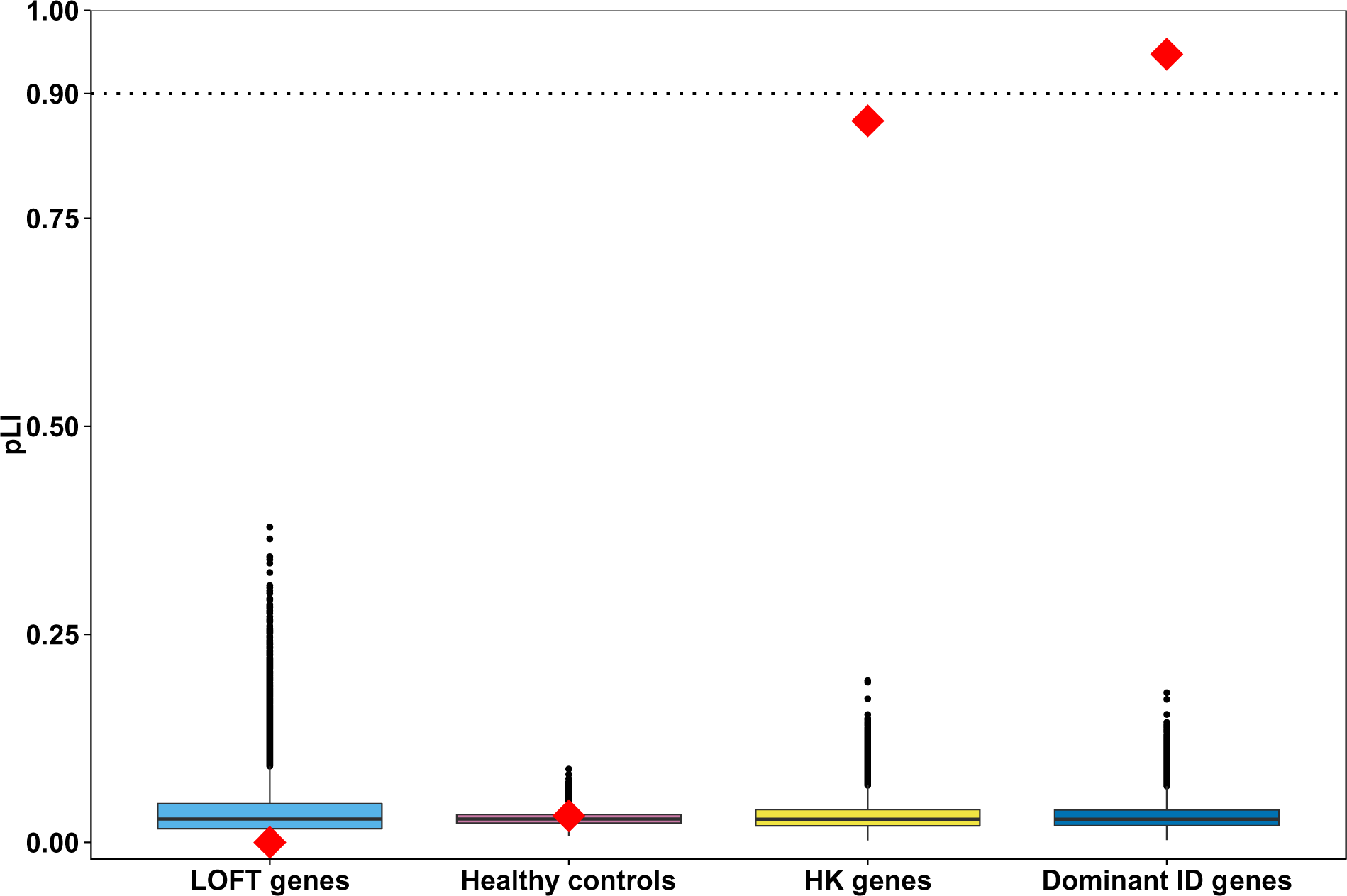
Gene set based evaluation of pLI. The median pLI median per gene set is depicted by a red diamond. Based on 100,000 simulations (**Online methods**) the median pLI for the loss-of-function tolerant (LOFT genes; N= 170) genes was significantly lower (light blue box; observed 9.33×10^−9^ vs. expected 0.03; empirical p-value: <1×10^−5^) than expected. For the healthy control set (Healthy controls; N=1,359) the observed median pLI matched the expected median pLI (pink box; observed 0.03 vs. expected 0.03; empirical p-value: 0.31). For the “house-keeping” (HK; N= 404; yellow box) and dominant ID (Dominant ID genes; N=444; dark blue box) gene sets the observed median pLI is significantly higher than the expected median pLI (Observed: 0.87 vs. expected: 0.03; empirical p-value <1×10^−5^; Observed: 0.95 vs. expected: 0.03; empirical p-value <1×10^−5^, respectively).

**Supplementary Figure 13.**
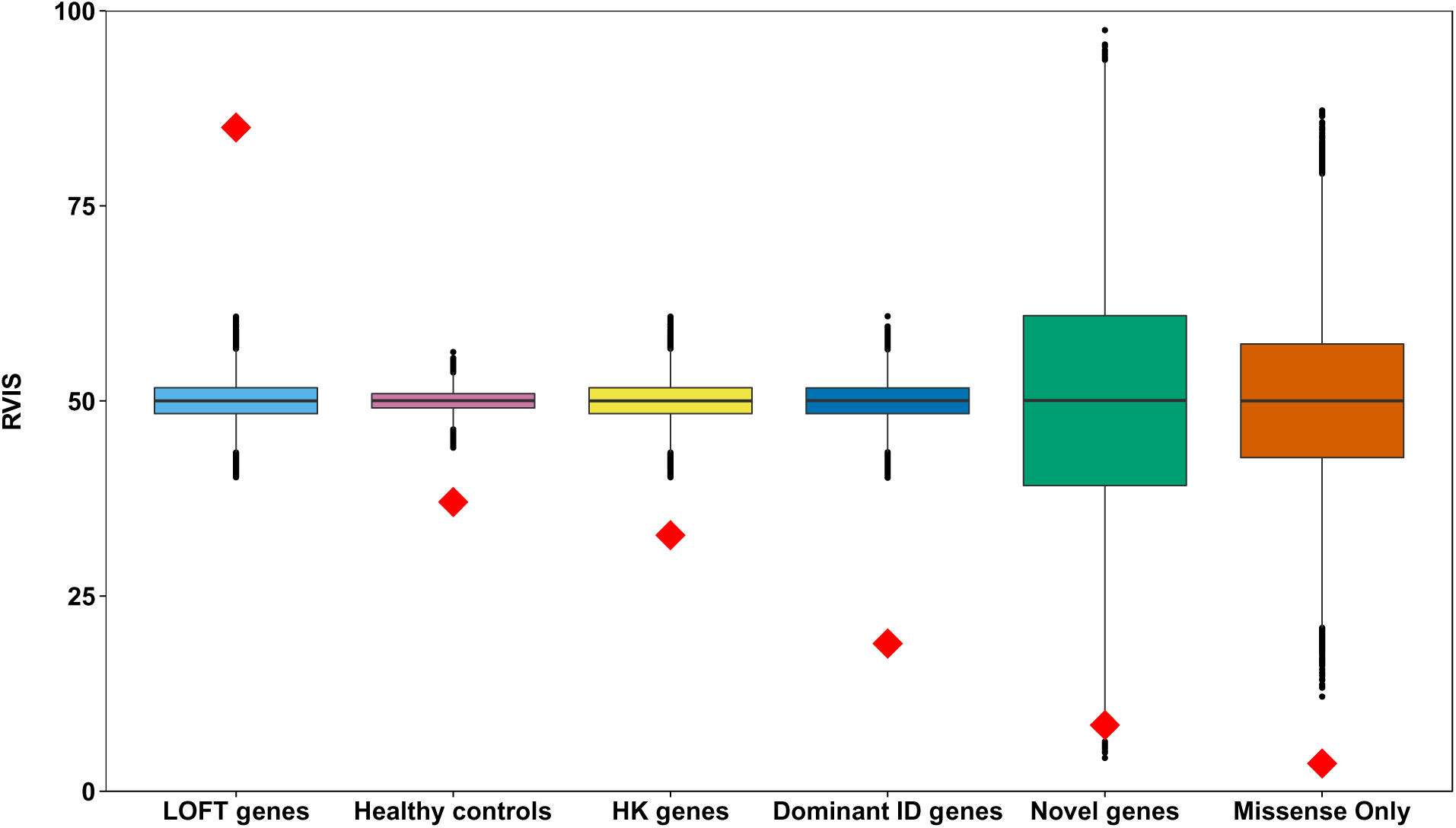
Gene set based evaluation of RVIS. The median RVIS median per gene set is depicted by a red diamond. Based on the simulations (**Online methods**) we identified a significant higher (light blue box; observed 85.04 vs. expected 50; empirical p-value: <1×10^−5^) median RVIS for the loss-of-function tolerant (LoFT) genes which is in line with the tolerant nature of this gene set. For the healthy control set the observed median RVIS was, to our surprise and unlike the pLI, significantly lower than the expected median RVIS (pink box; observed 37.05 vs. expected 50.00; empirical p-value: <1×10^−5^). For the “house-keeping” and dominant ID gene sets the observed median RVIS is significantly lower than the expected median RVIS (Yellow box; observed: 32.80 vs. expected: 50.00; empirical p-value <1×10^−5^; dark blue box observed: 18.92 vs. expected: 50.00; empirical p-value <1×10^−5^, respectively). The set of ten novel candidate ID genes has an observed median RVIS of 8.47 (green box; empirical p-value = 4.60x10^−4^). For the 21 dominant ‘missense only’ genes (with at least 3 missense mutations in the absence of LoF mutations) we observe the lowest median RVIS of 3.56 (orange box; empirical p-value <1×10^−5^) again illustrating that those known and novel candidate dominant ID genes that harbor only missense variants are among the most intolerant ID genes.

## References

1. Gulsuner, S., et al. Cell 154, 518–529 (2013).

2. Iossifov, I., et al. Nature 515, 216–221 (2014).

3. Gilissen, C., et al. Nature (2014).

4. O’Roak, B.J., et al. Nat Commun 5, 5595 (2014).

5. The Deciphering Developmental Disorders Study. Nature 519, 223–228 (2015).

6. Samocha, K.E., et al. Nat Genet 46, 944–950 (2014).

7. Luscan, A., et al. J Med Genet 51, 512–517 (2014).

8. Lumish, H.S., Wynn, J., Devinsky, O. & Chung, W.K. J Autism Dev Disord 45, 3764–3770 (2015).

9. Lek, M., et al. bioRxiv (2015).

10. Krumm, N., O’Roak, B.J., Shendure, J. & Eichler, E.E. Trends Neurosci 37, 95–105 (2014).

11. McRae, J.F., et al. bioRxiv (2016).

12. Neveling, K., et al. Hum Mutat 34, 1721–1726 (2013).

13. de Ligt, J., et al. N Engl J Med (2012).

14. Strom, S.P., et al. Genet Med 16, 510–515 (2014).

15. Gene panel intellectual disability. (Genome Diagnostics Nijmegen, Nijmegen, 2015).

16. Kong, A., et al. Nature 488, 471–475 (2012).

17. Goeman, J.J. & Solari, A. Stat Med 33, 1946–1978 (2014).

18. MacArthur, D.G., et al. Science 335, 823–828 (2012).

19. Zhu, J., He, F., Song, S., Wang, J. & Yu, J. BMC Genomics 9, 172 (2008).

20. Petrovski, S., Wang, Q., Heinzen, E.L., Allen, A.S. & Goldstein, D.B. PLoS Genet 9, e1003709 (2013).

## Supplementary References

1. Ozer, B.K. Econ Hum Biol 5, 280–301 (2007).

2. Gilissen, C., et al. Nature (2014).

3. Rauch, A., et al. Lancet 380, 1674–1682 (2012).

4. de Ligt, J., et al. N Engl J Med 367, 1921–1929 (2012).

5. The Deciphering Developmental Disorders Study. Nature 519, 223–228 (2015).

6. Neale, B.M., et al. Nature 485, 242–245 (2012).

7. Iossifov, I., et al. Nature 515, 216–221 (2014).

8. EuroEPINOMICS-RES Consortium; Epilepsy Phenome/Genome Project; Epi4K Consortium. Am J Hum Genet 95, 360–370 (2014).

9. Xu, B., et al. Nat Genet 43, 864–868 (2011).

10. McCarthy, S.E., et al. Mol Psychiatry 19, 652–658 (2014).

11. Gulsuner, S., et al. Cell 154, 518–529 (2013).

12. Xu, B., et al. Nat Genet 44, 1365–1369 (2012).

13. Fromer, M., et al. Nature 506, 179–184 (2014).

14. Hirunsatit, R., et al. Pharmacogenet Genomics 19, 53–65 (2009).

15. Carvill, G.L., et al. Am J Hum Genet 96, 808–815 (2015).

16. Babbs, C., et al. J Med Genet 51, 737–747 (2014).

17. Omer, C.A., Miller, P.J., Diehl, R.E. & Kral, A.M. Biochem Biophys Res Commun 256, 584–590 (1999).

18. Yochum, G.S., Sherrick, C.M., Macpartlin, M. & Goodman, R.H. Proc Natl Acad Sci U S A 107, 145–150 (2010).

19. Poy, F., Lepourcelet, M., Shivdasani, R.A. & Eck, M.J. Nat Struct Biol 8, 1053–1057 (2001).

20. Yazar, S., Gooden, G.E., Mackey, D.A. & Hewitt, A.W. PLoS One 9, e108490 (2014).

21. Lek, M., et al. bioRxiv (2015).

22. Iasevoli, F., Tomasetti, C. & de Bartolomeis, A. Neurochem Res 38, 1–22 (2013).

23. Zalfa, F., et al. Nat Neurosci 10, 578–587 (2007).

